# Dose-Dependent Softening of Bacterial Model Membranes by Structurally Distinct Antimicrobial Peptides: A Coarse-Grained Molecular Dynamics Study

**DOI:** 10.64898/2026.06.18.733300

**Authors:** Roni Saiba, Krishnakanth Baratam, Debayan Chakraborty, Satyavani Vemparala

## Abstract

Antimicrobial peptides (AMPs) act at the membrane interface, where they remodel lipid packing defects and redistribute lateral stresses, yet a quantitative, dose-dependent understanding of how they alter membrane mechanical properties remains incomplete. We use coarse-grained MARTINI 3 molecular dynamics simulations to systematically characterize the mechanical and microstructural response of a 70:30 POPE:POPG bilayer to three AMPs spanning distinct structural classes: aedesin (alpha-helical, 2MMM), arenicin-1 (beta-hairpin, 2JSB), and indolicidin (disordered, 1G89). For each peptide we vary the surface loading from one to four peptides per leaflet and extract the bending modulus *K*_*c*_, the area compressibility modulus *K*_*A*_, peptide localization depth, bilayer thickness, peptide-lipid and peptide-peptide spatial organization, and leaflet-resolved lipid packing defect distributions. All three peptides soften *K*_*c*_ monotonically with loading, but at per-peptide rates that span a threefold range and order systematically by structural class: − 1.39 ± 0.09, − 0.66 ± 0.04, and − 0.44 ± 0.01 k_B_T per peptide for aedesin, arenicin-1, and indolicidin, respectively. The tilt and twist moduli remain invariant across all conditions, indicating that the perturbation operates selectively on long-wavelength collective deformation modes. *K*_*A*_ softens for the two structured peptides but is statistically indistinguishable from the control for indolicidin, a dissociation we trace to a supraphosphate adsorption versus interfacial insertion dichotomy: structured peptides sit above the phosphate plane and act as supraphosphate wedges, while the disordered peptide threads into the interface without coherently displacing lipids. Independent geometric, spatial-organization, and microstructural observables corroborate this framework, with the deep versus shallow defect remodeling asymmetry providing a clean microstructural counterpart to the *K*_*c*_–*K*_*A*_ dichotomy. Acyl chain order parameters resolve the per-lipid splay from the bilayer-averaged response and show that the per-lipid perturbation tracks conformational state rather than peptide length: the two structured peptides impose comparable per-lipid chain disordering despite differing in length, while the disordered peptide imposes far less. These findings establish a quantitative connection between peptide-induced defect remodeling and the elastic response of the bilayer, and suggest a design principle in which conformational restriction maximizes the per-peptide membrane perturbation, motivating experimental tests on stapled-peptide AMP analogs.

## I. INTRODUCTION

Antimicrobial peptides (AMPs) are key components of the innate immune system across all domains of life, offering a broad-spectrum defense against bacterial pathogens [1, 2]. In the face of escalating antimicrobial resistance (AMR), which represents one of the most pressing global health challenges [3], AMPs have attracted sustained interest as potential alternatives to conventional antibiotics [4, 5]. Despite decades of research, however, a quantitative understanding of how AMPs alter the mechanical properties of target membranes remains incomplete.

The mechanical behavior of lipid bilayers is governed by a set of elastic moduli that encode the free energy cost of various deformation modes. Within the Helfrich-Canham-Evans continuum framework [6–8], the bending modulus K_*c*_ serves as the principal elastic constant characterizing the resistance of a membrane to curvature deformations. This quantity directly determines the energetics of membrane shape fluctuations, vesicle budding, pore formation, and the conformational dynamics of membrane-embedded proteins. Within this continuum description, K_*c*_ is intimately connected to the molecular-level distribution of internal stresses: as shown by Cantor [9], the bending modulus is determined by the second moment of the lateral pressure profile Π(*z*) across the bilayer thickness, establishing that any membrane-active agent that redistributes the lateral pressure—particularly in the acyl chain region—will necessarily alter K_*c*_. Measuring K_*c*_ and understanding how it is modulated by membrane-active agents is therefore of fundamental importance for elucidating antimicrobial mechanisms at the molecular level.

Experimentally, the bending modulus has been measured through techniques such as flicker spectroscopy [10, 11], micropipette aspiration [8, 12], and X-ray scattering from multilayer stacks [13]. Computationally, the traditional approach involves spectral analysis of bilayer undulations sampled from molecular dynamics (MD) trajectories, as formalized within the Helfrich framework [14]. However, this standard spectral method requires large membrane patches (typically exceeding 1000 lipids) to adequately sample long wavelength undulations. A significant methodological advance was introduced by Watson et al. [15], who demonstrated that the bending modulus can be extracted from fluctuations in lipid orientation vectors rather than membrane height, enabling reliable measurements from substantially smaller simulation systems of approximately 400-650 lipids. This lipid orientation approach was subsequently placed on rigorous theoretical footing by Levine et al. [16], who showed that the power spectra of lipid tilt vectors simultaneously yield the bending modulus K_*c*_, the tilt modulus K_*θ*_, and the twist modulus K_*tw*_ from fully atomistic simulations. The theoretical underpinnings of this coupling between lipid tilt and membrane curvature were further developed by Terzi and Deserno [17], who constructed a consistent quadratic curvature-tilt theory that accounts for splay-tilt coupling and lipid twist within a unified elastic framework. Other computational approaches have also been developed, including real-space fluctuations [18, 19] and density correlations. It is interesting to note that although different methods do not give the same values for different moduli, the trends remain similar across methods.

A separate but related aspect of AMP-membrane interactions involves lipid packing defects. These defects, which arise from thermal fluctuations in lipid headgroup packing, serve as critical sites for the initial binding and insertion of amphipathic molecules, including AMPs. The PackMem tool, developed by Gautier et al. [20], provides a systematic framework for identifying and classifying these defects as deep (penetrating below the glycerol level toward the bilayer center) or shallow (remaining above the glycerol reference plane). Prior work from our group [21, 22] has demonstrated that AMPs from different structural classes, namely alpha helical, beta sheet, alpha+beta, and disordered peptides, interact with and modulate these packing defects in distinct ways. However, the question of how multiple AMPs acting in concert, at varying surface concentrations, collectively influence membrane mechanical properties has remained largely unexplored at the molecular level. In biological settings, AMPs are expected to act at finite concentrations, and their effects on membrane structure may exhibit nonlinear dose response behavior arising from cooperative interactions, membrane saturation, or distinct binding configurations. Furthermore, the relationship between peptide induced changes in elastic moduli and the concurrent modulation of lipid packing defect characteristics has not been systematically examined.

The question of how packing defects relate to membrane mechanics acquires additional significance in light of a long-standing debate in membrane biophysics: whether membrane-active amphipathic molecules sense the geometric curvature of the membrane, or whether they sense the lipid packing defects that are enhanced at curved surfaces. The seminal work of Bigay, Antonny, and coworkers on the ALPS motif established that certain amphipathic helices do not recognize curved membrane geometry per se; rather, they recognize the packing defects that arise from the mismatch between membrane curvature and lipid geometry [23, 24]. Subsequent computational work by Vanni et al. showed that the correlation between amphipathic helix partitioning and packing defect distributions quantitatively explains how both curvature and lipid composition modulate protein recruitment, with these two factors acting cumulatively [25, 26]. Atomistic simulations further revealed that packing defects on curved membranes increase not only in number but also in a nontrivial way, with their size distribution broadening on more highly curved surfaces [27]. An alternative but complementary viewpoint, advanced by Campelo and Kozlov, argued that the relevant quantity sensed by shallow insertions is the intra-membrane stress rather than curvature itself, and that the binding affinity is determined by the resultant stress independently of how it was generated [28]. Recent work by van Hilten et al. has further shown that deeply inserted amphipathic helices can sense negative curvature, a regime where packing defects are suppressed, suggesting that the full depth-resolved lateral pressure profile, rather than surface-level defects alone, governs the sensing response [29]. In our earlier works [22, 30, 31], we have shown the role of lipid packing defects in partitioning of membrane active agents into the model membranes and also role functional groups as well as secondary structures on generation as well as detection of lipid packing defects

The crucial point connecting these perspectives to the present study is that packing defects, internal membrane stresses, and elastic moduli are not independent quantities but they are different manifestations of the same underlying lateral pressure profile. Packing defects are a surface manifestation of the internal stress distribution; the elastic moduli are its integral moments. However, protein-induced remodeling of packing defects altering the mechanical properties that govern membrane curvature has remained unexplored. The present work addresses this reverse direction in the context of antimicrobial peptides. In this work, we address these questions using coarse-grained molecular dynamics simulations with the MARTINI 3 force field [32], which affords access to the microsecond timescales and large system sizes required for reliable elastic modulus measurements in the presence of peptide inclusions. We systematically vary the number of bound AMPs from one to four per membrane for three representative AMPs spanning distinct structural classes: 2MMM (Aedesin), an alphahelical peptide; 2JSB (Arenicin-1), a beta-hairpin peptide; and 1G89 (Indolicidin), a disordered cationic peptide. For each system we quantify the resulting changes in the bending modulus K_*c*_, the area compressibility modulus K_*A*_, the peptide localization depth relative to the phosphate plane, the bilayer thickness distribution, the peptide-peptide and peptide-lipid spatial organization, and the leaflet-resolved lipid packing defect distributions. Multiple independent replicas are performed for each condition to capture the effects of metastable peptide binding configurations. Our results reveal a clear dose dependent softening of the membrane bending modulus that depends systematically on the structural class of the AMP, accompanied by a structural class specific softening of the area compressibility modulus that dissociates for the disordered peptide. We rationalize these differential mechanical responses through a supraphosphate adsorption versus interfacial insertion framework in which the structured peptides act as supra-headgroup wedges while the disordered peptide threads into the interface without coherently displacing lipids. Independent geometric, spatial, and microstructural observables corroborate this framework and additionally reveal a structural class specific short-range peptide-peptide correlation that emerges for the alpha-helical AMP at the highest concentration studied. Together, these findings establish a quantitative framework connecting AMP structural identity, surface concentration, and the resulting perturbation of membrane mechanical and interfacial properties.

## II. MATERIALS AND METHODS

### A. System construction

All simulations were done using the MARTINI 3 coarse grained force field [32, 33]. A model bacterial membrane was constructed using the Martini Maker module of CHARMM-GUI [34, 35], consisting of 1020 lipids distributed equally between two leaflets (510 per leaflet). Each leaflet contained 357 POPE and 153 POPG lipids, yielding a 70:30 POPE:POPG molar ratio that approximates the inner membrane composition of Gram-negative bacteria. The system was solvated with MARTINI coarse-grained water and neutralized with the appropriate number of counterions, maintaining a physiological salt concentration of 150 mM NaCl.

Three representative antimicrobial peptides spanning distinct structural classes were selected based on our earlier classification study [21] of AMPs from the DRAMP database [36]: 2MMM (aedesin) [37], a 36-residue alpha-helical peptide (sequence GGLKKLGKKL-EGAGKRVFKASEKALPVVVGIKAIGK, net charge +8); 2JSB (arenicin-1) [38], a 21-residue beta-hairpin peptide stabilized by a Cys3–Cys20 disulfide bond (sequence RWCVYAYVRVRGVLVRYRRCW, net charge +6); and 1G89 (indolicidin) [39, 40], a 13-residue disordered cationic peptide (sequence ILPWKWPWWP-WRR, net charge +4, C-terminal amide). Coarse-grained representations of the structured peptides (aedesin, arenicin-1) were generated using the ElNeDyn elastic network approach [41], which preserves secondary and tertiary structure through harmonic backbone restraints. For indolicidin, the recently developed Martini3-IDP parameterization [42] was used, which omits the elastic network and treats the peptide as fully disordered to enable sampling of conformational flexibility. For each peptide type, systems were prepared with n = 1, 2, 3, and 4 peptides placed in the aqueous phase above the membrane surface, together with a lipid-only control system. The peptides were positioned in the solution phase at a sufficient distance from the membrane to avoid initial steric contacts, and binding to the membrane proceeded spontaneously during the simulation. A representative image of the final configurations observed in each system, shown for n = 4 for each peptide, is presented in Figure 1. All configurations were visualised using VMD [43], preprocessed using the MartiniGlass framework [44].

**FIG. 1.**
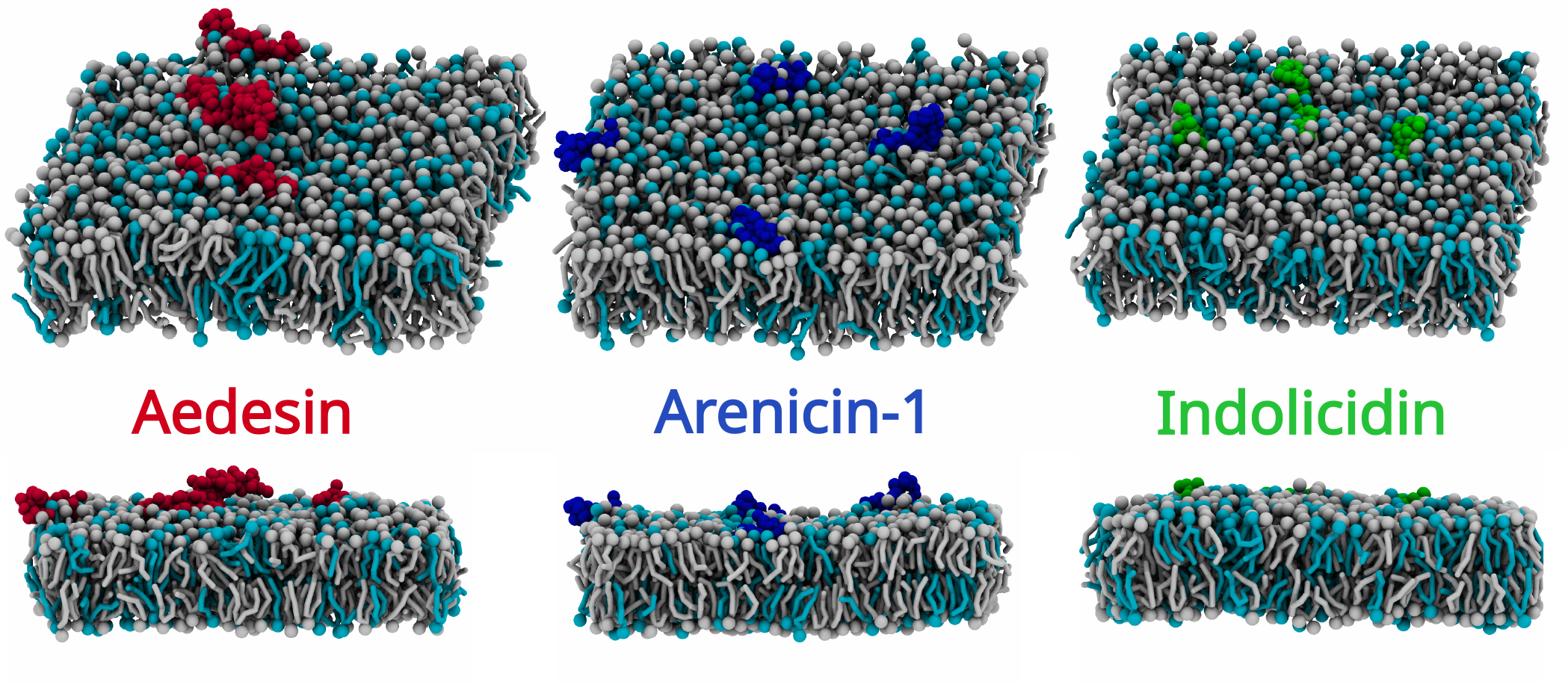
The three systems of interest in this study: For the purpose of illustration, we have chosen the final configuration observed in one of the replicas containing 4 peptides. The lipids are shown in grey(POPE) and cyan(POPG) colours with the phosphate head group and lipid tail groups represented using bead and licorice format, respectively. The peptides are represented using red, blue and green beads for Adesein, Arenicin-1 and Indolicidin, respectively. The water and ion beads were turned off during visualisation for clarity.

### B. Simulation protocol

All simulations were performed using GROMACS 2025.3 [45]. Nonbonded interactions followed the standard Martini 3 parameters: reaction-field electrostatics [46] with a relative dielectric constant of 15 and a Coulomb cutoff of 1.1 nm, and a van der Waals cutoff of 1.1 nm with the potential-shift-Verlet modifier. The Verlet neighbor list scheme was used with a list update frequency of 20 steps and a buffer tolerance of 0.005 kJ mol^−1^ ps^−1^. Periodic boundary conditions were applied in all three dimensions. Each system was energy minimized using steepest descent, followed by a five-step equilibration protocol in which position restraints on peptide backbone beads and lipid headgroup beads along the membrane normal were progressively reduced over approximately 4.75 ns, while the integration timestep was gradually increased from 2 fs to 20 fs. The lipid-only control employed an additional initial soft-core minimization step with extended cutoffs to resolve steric clashes, following the CHARMM-GUI [35, 47, 48] protocol for Martini simulations. Complete details of the restraint force constants, timesteps, and barostat choices for each equilibration stage are provided in Supplementary Table SX.

Production simulations were run for 1.0 microsecond each (50,000,000 steps, dt = 20 fs) with the Parrinello-Rahman barostat [49] under semi-isotropic coupling (*τ*_*p*_ = 12.0 ps, compressibility 3 × 10^−4^ bar^−1^, reference pressure 1.0 bar) and the v-rescale thermostat [50] at 3×10 K (*τ*_*T*_ = 1.0 ps). Temperature coupling was applied separately to three groups (protein, membrane, and solute) for peptide-containing systems and two groups (membrane, and solute) for the lipid-only control. Trajectory coordinates were saved every 100 ps. Independent repli-cas were generated by assigning distinct random velocity seeds at the start of each production run, thereby sampling independent regions of phase space from the same equilibrated configuration. Multiple independent replicas were performed for each peptide concentration condition (N ranging from 6 to 14 per condition) and five for the lipid-only control.

### C. Analysis of membrane elastic moduli

Membrane elastic moduli were extracted using the Watson-Levine spectral analysis [15, 16] of lipid orientation fluctuations. For each lipid, an orientation vector was defined from the midpoint of the headgroup beads (PO4 and GL2) to the midpoint of the terminal tail beads (C4A and C4B), following the coarse-grained analogue of the definition employed by Levine et al. [16]. These vectors were assigned to an M × M grid (M = 14) spanning the lateral box dimensions, with separate grids for each leaflet. Empty grid cells were filled by weighted averaging over periodic nearest neighbors. The discrete Fourier transform of the gridded orientation field was computed for each frame, and the power spectra of the longitudinal and transverse tilt components were accumulated.

The bending modulus K_*c*_ was obtained from the longitudinal spectrum in the low-wavevector regime (q ≤ 1.0 nm^−1^), the tilt modulus K_*θ*_ from both spectra in the high-wavevector regime (0.3 ≤ q ≤ 2.5 nm^−1^), and the twist modulus K_*tw*_ from the transverse spectrum where the difference between longitudinal and transverse components isolates the twist contribution. The analysis was performed over the final 500 ns of each trajectory, sampled at an effective interval of 500 ps, yielding 1000 frames per replica. The initial 500 ns was discarded to allow for peptide binding and equilibration of the peptide-membrane complex. Errors on individual replicas were estimated from spectral fitting uncertainty; the inter-replica standard deviation, which exceeded within-replica errors for most conditions, is reported as the primary uncertainty measure.

The dose dependence of K_*c*_ on peptide number was characterized by weighted least-squares (WLS) regression of the per-replica K_*c*_ values on peptide concentration, with weights proportional to the inverse variance of the inter-replica K_*c*_ distribution at each concentration to account for heteroscedasticity. The regression was restricted to n ≥ 1, since the n = 0 *→* 1 transition conflates initial peptide binding with concentration-dependent effects. Slope p-values were assessed against the null hypothesis of zero softening.

As a microscopic probe of acyl chain ordering, we computed the coarse-grained order parameter 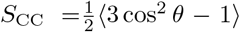 for each consecutive pair of tail beads, where *θ* is the angle between the bead-bead bond vector and the bilayer normal, yielding three values along each of the sn-1 (unsaturated) and sn-2 (saturated) chains. Order parameters were accumulated separately for the upper (peptide-facing) and lower leaflets. To distinguish the bilayer-averaged response from the response of lipids in direct contact with the peptide, two scopes were evaluated at each concentration: a global scope averaging over all lipids in the leaflet, and a local scope restricted to lipids whose headgroup lay within 2.5 nm laterally of the nearest peptide center of mass, using the same first-shell neighbor definition employed for the localization analysis. The lipid-only control provided the n = 0 reference, and the perturbation is reported as Δ*S*_CC_ (peptide minus control) per bead position.

### D. Analysis of lipid packing defects

Lipid packing defects were analyzed using the PackMem software developed by Gautier et al. [20]. MARTINI 3-compatible parameter files for PackMem were generated in-house, following the recommendations of van Hilten et al. [51] for adapting the tool to coarsegrained force fields. Classification as deep or shallow was based on penetration depth relative to the glycerol-level bead (GL2 in the MARTINI mapping). For each frame, the number of defect sites, individual defect areas, and total defect area were computed separately for the upper and lower leaflets. The analysis was performed over the final 300 ns of each trajectory (3000 frames at 100 ps interval). Frame-level statistics were pooled across all replicas for each condition, with the lipid-only control serving as the shared reference for all peptide systems.

## III. RESULTS

### A. Dose-dependent change of membrane bending modulus

To establish the baseline mechanical properties of the model bacterial membrane, we first analyzed the lipid-only control system. Using the Watson-Levine spectral analysis [15, 16] of lipid orientation fluctuations with M = 14 grid divisions and a wavevector cutoff of q_*max*_ = 1.0 nm^−1^, we obtained a bending modulus of K_*c*_ = 16.47 ± 0.27 k_B_T (mean ± SD across five independent replicas). The grid divisions were chosen such that the area of each grid cell is close to the value recommended by Watson and Levine. This K_*c*_ value is consistent with the expected range for POPE-containing membrane models using the MARTINI 3 force field. The tilt modulus was measured as K_*θ*_ = 26.257 ± 0.186 k_B_T, and the twist modulus as K_*tw*_ = 1.907 ± 0.062 k_B_T, both exhibiting low inter-replica variance and serving as reference values for evaluating peptide-induced perturbations.

In contrast to the control, all three peptide systems exhibit a clear monotonic decrease in K_*c*_ with increasing peptide concentration, as shown in Figure 2. To quantify the dose-dependent softening, we performed a weighted least-squares (WLS) linear regression of K_*c*_ on peptide number over the range n = 1 to n = 4. The fit yields a softening rate that depends strongly on the structural class of the peptide.

**FIG. 2.**
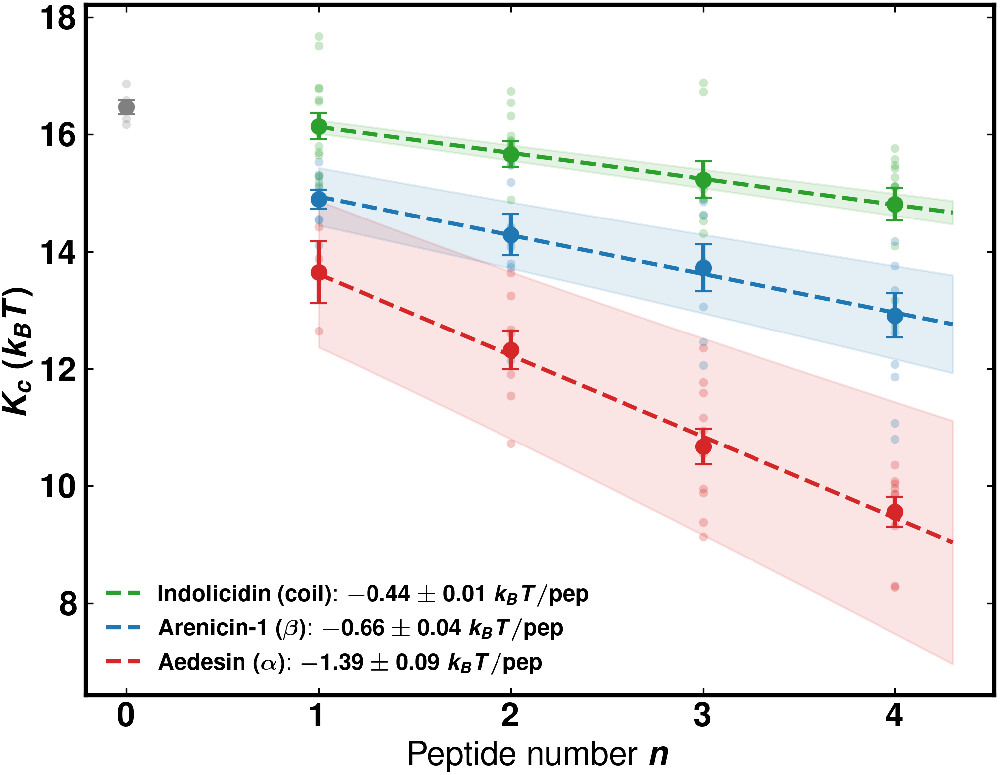
Membrane bending modulus K_*c*_ as a function of peptide number for the three AMP systems studied. Light symbols show individual replica values, large filled symbols show the per-concentration means with standard error, and the dashed line is the weighted least-squares fit over n ≥ 1 with the 95% confidence interval shown as a shaded band. The softening rate increases systematically with peptide structural order: -0.44 ± 0.01 k_B_T/peptide for indolicidin, -0.66 ± 0.04 k_B_T/peptide for arenicin-1, and -1.39 ± 0.09 k_B_T/peptide for aedesin. Peptide number, n=0 shows control simulations.

For indolicidin, K_*c*_ decreased from 16.14 ± 0.23 k_B_T at n = 1 to 14.81 ± 0.28 k_B_T at n = 4 (uncertainties reported as the standard error of the mean across replicas), corresponding to a 10.0% reduction relative to the control. The WLS slope was −0.44 ± 0.01 k_B_T/peptide (p = 0.001). For arenicin-1, the softening was substantially stronger: K_*c*_ dropped from 14.90 ± 0.16 k_B_T at n = 1 to 12.91 ± 0.38 k_B_T at n = 4, a 21.6% reduction, with a WLS slope of −0.66 ± 0.04 k_B_T/peptide (p = 0.003). For aedesin, the effect was strongest, with K_*c*_ decreasing from 13.65 ± 0.53 k_B_T at n = 1 to 9.56 ± 0.26 k_B_T at n = 4, a 41.9% reduction. The WLS slope was −1.39 ± 0.09 k_B_T/peptide (p = 0.004), more than three times that of indolicidin.

The threefold range in softening rates across structural classes points to a systematic dependence of the membrane mechanical response on peptide architecture, with structured peptides perturbing the bilayer more efficiently than their disordered counterpart on a per-molecule basis. The alpha-helical aedesin produces the largest effect, the beta-hairpin arenicin-1 produces an intermediate one, and the disordered indolicidin, which lacks a defined amphipathic surface and instead samples a broad conformational ensemble at the membrane interface, produces the mildest perturbation. The geometric origin of this structural-class ordering, including the relative depth of peptide localization with respect to the lipid headgroups and its consequences for the mode of mechanical coupling, is established in Section III C. The inter-replica variance in K_*c*_ also grows with peptide number for all three systems, reflecting the increasing diversity of accessible metastable peptide-membrane configurations at higher surface concentrations.

In marked contrast to the bending modulus, the tilt modulus K_*θ*_ remained essentially invariant across all concentrations and peptide types, fluctuating within a narrow range of 26.2 ± 0.2 k_B_T, with no systematic trend discernible above the statistical uncertainty. Similarly, the twist modulus K_*tw*_ showed no significant concentration dependence, remaining in the range of 1.89 ± 0.06 k_B_T for all three systems. The constancy of K_*θ*_ and K_*tw*_ indicates that the peptide-induced perturbation operates selectively on the long-wavelength collective deformation modes captured by the bending modulus, while leaving the local orientational elasticity of individual lipid molecules effectively unperturbed.

### B. Dose-dependent change of area compressibility modulus

Beyond the bending response, the area compressibility modulus K_*A*_ provides an independent measure of bilayer mechanical perturbation, capturing the in-plane elastic resistance against isotropic area dilation rather than the long-wavelength out-of-plane bending. K_*A*_ was obtained from box-area fluctuations in the NPT trajectories via the standard relation K_*A*_ = k_B_T ⟨*A*⟩ /Var(A) [52], evaluated over the same production window used for the spectral bending modulus analysis.

For the lipid-only control, the area compressibility modulus was K_*A*_ = 204.0 ± 1.8 mN/m (mean ± SEM across five independent replicas). At n = 1, all three peptide systems yielded K_*A*_ values in a narrow range of 203–205 mN/m (indolicidin: 204.8 ± 2.6 mN/m, arenicin-1: 204.9 ± 2.2 mN/m, aedesin: 202.7 ± 6.6 mN/m), consistent with the control value and internally consistent with the bending modulus through the polymer brush relation K_*c*_ = K_*A*_ t^2^/48 [12] for an effective bilayer thickness of approximately 3.9 nm, providing a cross-check on the two independent elastic moduli measurements. With increasing peptide concentration, the three peptides exhibited markedly divergent K_*A*_ behavior, in contrast to the qualitatively universal K_*c*_ softening reported in Section III A (Figure 3).

**FIG. 3.**
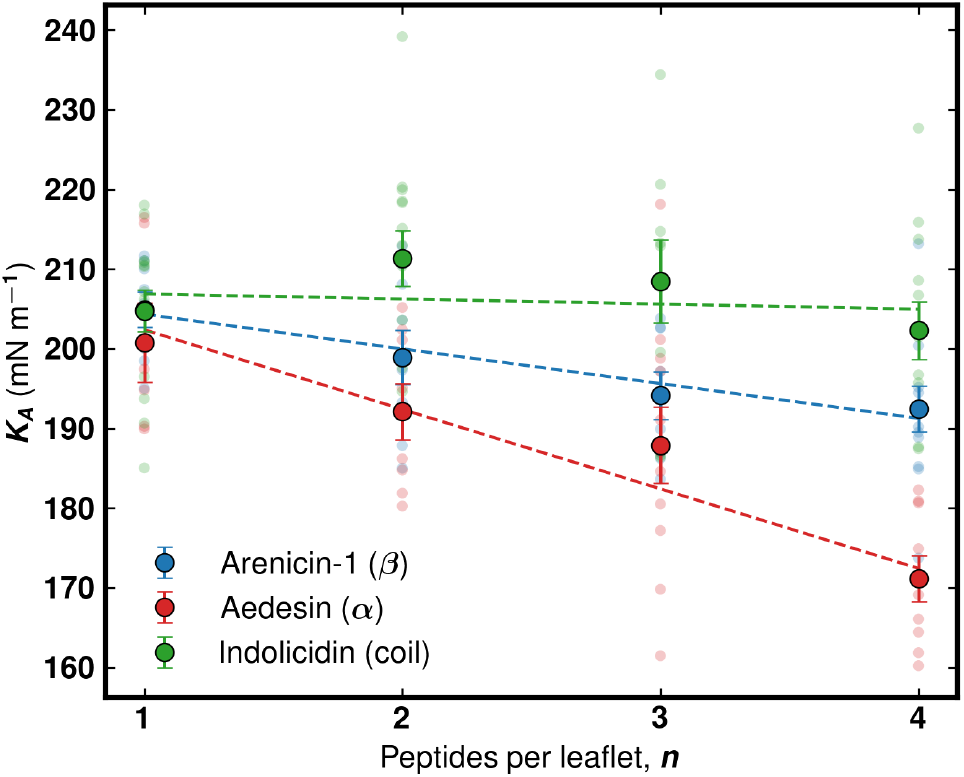
Area compressibility modulus K_*A*_ as a function of peptide number for the three AMP systems. Light symbols show individual replica values, large filled symbols show per-concentration means with standard error of the mean, and dashed lines show weighted least-squares fits over n = 1 to n = 4. Aedesin and arenicin-1 exhibit clear dose dependent softening with WLS slopes of -8.78 ± 1.88 and -4.03 ± 0.92 mN/m per peptide respectively, while indolicidin shows no resolvable concentration dependence (slope -0.64 ± 1.93 mN/m per peptide).

For aedesin, K_*A*_ decreased monotonically from 202.7 ± 6.6 mN/m at n = 1 to 171.2 ± 2.9 mN/m at n = 4, a 15.5% reduction relative to the n = 1 value. The WLS slope was −8.78 ± 1.88 mN/m per peptide (p < 10^−4^). For arenicin-1, the softening was substantially milder: K_*A*_ decreased from 204.9 ± 2.2 mN/m at n = 1 to 193.7 ± 2.9 mN/m at n = 4 (slope −4.03 ± 0.92 mN/m per peptide, p < 10^−4^), a 5.5% reduction. In striking contrast, indolicidin showed no resolvable K_*A*_ dependence on concentration: the mean K_*A*_ remained in the range of 202–211 mN/m across all four concentrations, with a WLS slope of − 0.64 ± 1.93 mN/m per peptide (p = 0.74) that was statistically indistinguishable from zero.

This dissociation between the K_*c*_ and K_*A*_ responses for indolicidin is mechanistically informative. The two elastic moduli, although both global descriptors of the bilayer, are dominated by different regions of the bilayer cross-section. The bending modulus K_*c*_ quantifies the elastic cost of long-wavelength out-of-plane deformations and is sensitive to perturbations distributed across the entire bilayer thickness, with substantial contributions from peripheral lipid tilt and curvature stress in the headgroup region. The area compressibility modulus K_*A*_, by contrast, is dominated by the integrated chain-packing rigidity at the bilayer core, set by the entropic repulsion of densely packed acyl chains. A peptide that perturbs the interfacial mechanics without coherently displacing lipids in the membrane plane will therefore soften K_*c*_ but leave K_*A*_ largely unaffected. The observation that all three structural classes soften K_*c*_ but only the structured peptides soften K_*A*_ implies that the mode of peptide-lipid coupling differs qualitatively between the disordered and structured AMPs, despite all three peptides operating at the membrane surface. The geometric origin of this difference is established in the following section.

### C. Geometric origin of differential mechanical coupling

To rationalize the structural-class dependence of the K_*c*_ and K_*A*_ softening, we computed the time-averaged center-of-mass position of each peptide relative to the local phosphate (PO_4_) plane of its binding leaflet. Position was reported as the signed difference between the mean absolute z-distance of the peptide center of mass and the mean absolute z-distance of the PO_4_ headgroups, both referenced to the bilayer midplane, such that positive values indicate the peptide center of mass lies above the PO_4_ plane (i.e., in the headgroup-water region) and negative values indicate it lies below (i.e., inserted into the acyl chain region).

The three peptides occupied clearly distinguishable position regimes (Figure 5a). Indolicidin sat essentially flush with the PO_4_ plane, with a mean position of −0.08 ± 0.04 nm averaged across concentrations, consistent with the disordered backbone integrating into the interface while the tryptophan side chains thread marginally below into the upper acyl chain region [40]. Arenicin-1 sat at +0.22 ± 0.05 nm above the PO_4_ plane, consistent with its compact beta-hairpin structure resting within the headgroup region. Aedesin sat at +0.49 ± 0.07 nm above the PO_4_ plane, the highest of the three peptides, reflecting the inability of its extended, elastic-network-constrained alpha-helical structure to deform and embed deeper into the interface; the peptide instead remains perched on top of the headgroups within the headgroup-water region.

A finer-grained view of peptide localization is provided by the per-bead z-distributions relative to the peptide-facing PO_4_ plane (Figure 4), which separate backbone (BB) and sidechain (SC) contributions. At n = 4, the fraction of peptide beads residing below the PO_4_ plane is 0.15 for aedesin (BB 0.14, SC 0.17), 0.33 for arenicin-1 (BB 0.26, SC 0.36), and 0.56 for indolicidin (BB 0.39, SC 0.62). The disordered indolicidin therefore samples below-PO_4_ space with more than half of its beads, dominated by the deep penetration of its tryptophan and arginine side chains, despite a center-of-mass position essentially flush with the PO_4_ plane. The structured peptides keep the majority of their mass above PO_4_, with aedesin in particular retaining roughly 85% of its beads in the headgroup-water region. Notably, this ordering is the reverse of the peptide chain lengths: the longest peptide (aedesin, 36 residues) penetrates least and the shortest (indolicidin, 13 residues) penetrates most, establishing at the outset that below-PO_4_ occupancy is governed by conformational state rather than by the number of residues. The qualitative difference in how each peptide occupies the headgroup region, beyond the COM coordinate alone, underpins the differential defect remodeling reported in Section III G.

**FIG. 4.**
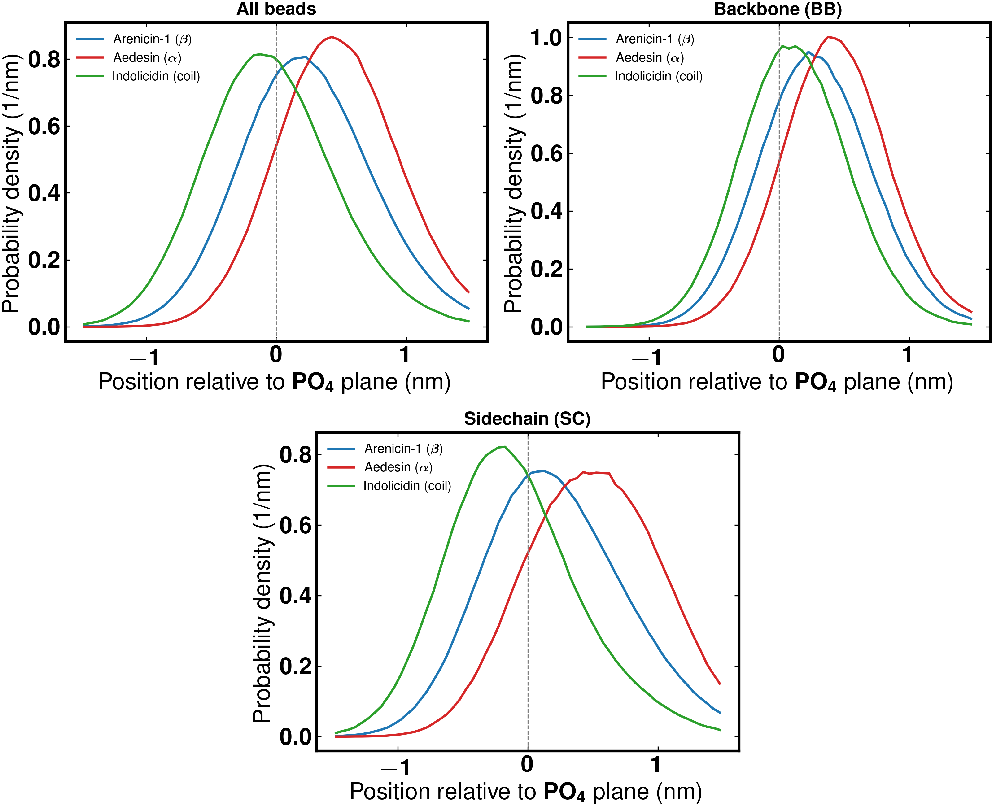
Probability density of peptide bead positions relative to the peptide facing PO_4_ plane at n = 4, for all beads (left), backbone beads only (center), and sidechain beads only (right). The vertical dashed line marks the PO_4_ plane; positive positions lie in the headgroup-water region and negative positions in the acyl chain region. The structured peptides (aedesin, arenicin-1) keep the bulk of both backbone and sidechain mass above PO_4_, whereas indolicidin places its backbone near PO_4_ and threads its sidechains below it, sampling below-PO_4_ space with the majority of its beads despite being the shortest peptide.

Two further features are evident from Figure 5a. First, the mean position is essentially invariant with peptide number for all three systems, demonstrating that the membrane binding mode saturates at n = 1 and that the dose-dependent moduli softening reported in Sections III A and III B arises from cumulative independent perturbation by surface-bound peptides rather than from concentration-dependent reorientation or progressive insertion. Second, the height-above-PO_4_ ordering of the three peptides directly tracks the magnitude of mechanical perturbation: the peptide producing the largest moduli softening (aedesin) sits farthest above the headgroup plane, while the peptide producing the smallest softening (indolicidin) sits closest to and marginally below it. A naive insertion-depth picture, which would predict softening to scale with penetration into the acyl chain region, is therefore inverted: the relevant geometric descriptor is the height of the peptide center of mass above the PO_4_ plane, set by the structural class of the peptide and the volume and lateral footprint of its body residing in the headgroup-water region.

**FIG. 5.**
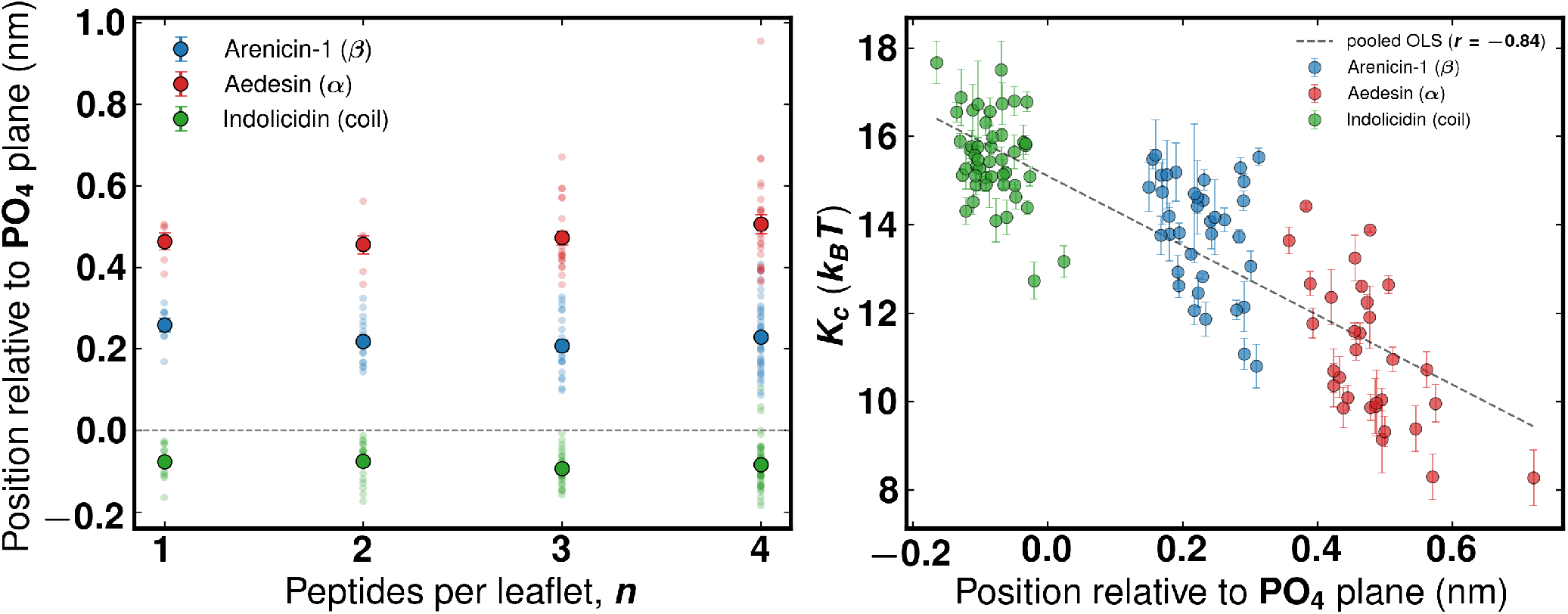
(a) Peptide center of mass position relative to the phosphate (PO_4_) plane as a function of peptide number. Positive values indicate localization above PO_4_ (in the headgroup-water region); negative values indicate insertion below PO_4_ (toward the acyl chain region). Light symbols show per-replica per peptide positions; large filled symbols show per-concentration means with standard error of the mean. The position is essentially invariant with n for all three peptides. (b) Per-replica spectral K_*c*_ plotted against the corresponding replica-averaged position, pooled across all concentrations; error bars are the spectral fit uncertainty in K_*c*_ for each replica. The dashed line is the pooled ordinary least-squares reference fit (Pearson *r* = −0.84). The three peptides occupy clearly separated regions of position space, with the height above the PO_4_ plane tracking the magnitude of K_*c*_ softening at the class level.

We tested the relationship between per-replica peptide localization and per-replica mechanical response by plotting the spectral K_*c*_ value from each replica against the corresponding replica-averaged peptide position (Figure 5b). Pooled across all three peptides and all concentrations, the two quantities exhibited a strong negative correlation (Pearson *r* = − 0.84, *p <* 10^−30^, *N* = 116): peptides positioned higher above the PO_4_ plane systematically associate with lower K_*c*_ (greater softening). Per-peptide correlations within each structural class were substantially weaker (arenicin-1 *r* = − 0.36, *p* = 0.03, *N* = 37; aedesin *r* = − 0.62, *p <* 10^−4^, *N* = 31; indolicidin *r* = − 0.38, *p <* 0.01, *N* = 48), confirming that the pooled correlation is driven primarily by the clean separation of the three peptide systems into distinct neighborhoods of position space rather than by a continuous trend within a single structural class. Within each peptide group, the position varies by less than 0.2 nm whereas the per-replica K_*c*_ varies by 4–5 k_B_T, with the latter dominated by configurational sampling at fixed n. The pooled correlation should therefore be interpreted as a class-level statement: peptide structural class, encoded geometrically by the height of the peptide center of mass above the PO_4_ plane, predicts the magnitude of bending modulus softening.

This geometric pattern, taken together with the dose-dependent K_*A*_ response in Section III B, is consistent with a supraphosphate adsorption versus interfacial insertion dichotomy for AMP-membrane mechanical coupling. The structured peptides, constrained by elastic network restraints to their native fold, sit on top of the headgroup region and act as supraphosphate wedges. The volume occupied by the rigid peptide backbone above the PO_4_ plane forces the underlying acyl chains to splay outward to compensate for the local displacement of headgroup material, producing both a strong perturbation of the long-wavelength bending response (K_*c*_ softens) and a measurable reduction of the chain-packing rigidity in the plane (K_*A*_ softens). The larger the supra-headgroup wedge, the greater the perturbation. This explains why aedesin, which sits highest above PO_4_ and presents the most extended amphipathic body to the interface, produces the largest effect on both moduli, and why arenicin-1, whose compact beta-hairpin rests at an intermediate height, produces an intermediate effect. The controlling variable is the supraphosphate wedge geometry set by the conformational state of the peptide, not its chain length: the height-above-PO_4_ and below-PO_4_ occupancy orderings both run opposite to the residue-count ordering (aedesin is the longest peptide yet penetrates least, indolicidin the shortest yet penetrates most), so a simple per-residue scaling of the perturbation is excluded by the localization data alone. Indolicidin, lacking secondary structure, adapts to the interface; its backbone lies flush along the PO_4_ plane and the tryptophan side chains thread into the interstitial space between lipid headgroups without coherently displacing them. This interfacial insertion mode disorders local lipid tilt and curvature stress at the interface, producing a measurable but modest K_*c*_ softening, but does not generate the coherent lipid displacement required to soften K_*A*_. The dissociation between the K_*c*_ and K_*A*_ responses for indolicidin is therefore not a paradox but a direct mechanical signature of conformational disorder at the membrane interface.

A quantitative consistency check on this picture is provided by the polymer brush relation [12], which connects the bending and area compressibility moduli through an effective bilayer thickness, 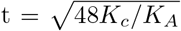. Combining the K_*c*_ values from Section III A with the K_*A*_ values from Section III B, the effective thickness at n = 1 is 4.02 nm for indolicidin, 3.86 nm for arenicin-1, and 3.72 nm for aedesin, indicating that even at the lowest concentration the structured peptides already produce a measurable reduction of the effective thickness relative to the disordered case. At n = 4, the effective thickness drops to 3.88 nm for indolicidin (a 3.5% reduction relative to its n = 1 value), 3.70 nm for arenicin-1 (4.1%), and 3.39 nm for aedesin (8.9%). The magnitude of effective thinning thus tracks the supraphosphate adsorption versus interfacial insertion hierarchy quantitatively: peptides that sit highest above the PO_4_ plane and possess the largest lateral footprint produce the strongest effective thinning of the bilayer core, while the disordered peptide that integrates into the interface produces only a modest effective thinning that arises almost entirely from the K_*c*_ reduction rather than from any K_*A*_ effect. This collapse of two independent elastic moduli into a single thickness coordinate provides a compact scalar descriptor of dose-dependent membrane perturbation by AMPs.

It should be noted that the strongly surface-adsorbed configurations of the structured peptides reflect, in part, the elastic network restraints employed in the MARTINI 3 representation, which preserve the native secondary structure and prevent the partial unfolding and chain insertion events observed for aedesin in atomistic simulations [21]. The analysis presented here therefore characterizes the mechanical coupling in the surface-adsorbed regime relevant to the early stages of AMP-membrane interaction, prior to any insertion or pore-forming events that may occur on longer timescales and through different molecular mechanisms.

### D. Acyl chain order parameters and the length independence of the wedge mechanism

The supraphosphate wedge mechanism makes a direct microscopic prediction: peptides that sit above PO_4_ and displace headgroup material should force the underlying acyl chains to splay, reducing the chain order parameter *S*_CC_ of the lipids in their footprint. Because the three peptides differ in chain length as well as in conformational state, the order parameter response also provides the most direct available test of whether the mechanical hierarchy is a length effect in disguise. We address this by separating the bilayer-averaged (global) chain-order response, which is extensive and scales with the number of lipids a peptide contacts, from the per-lipid (local) response of the lipids in direct contact with the peptide, which is intensive and reports the splay imposed on each footprint lipid (Figure 6).

**FIG. 6.**
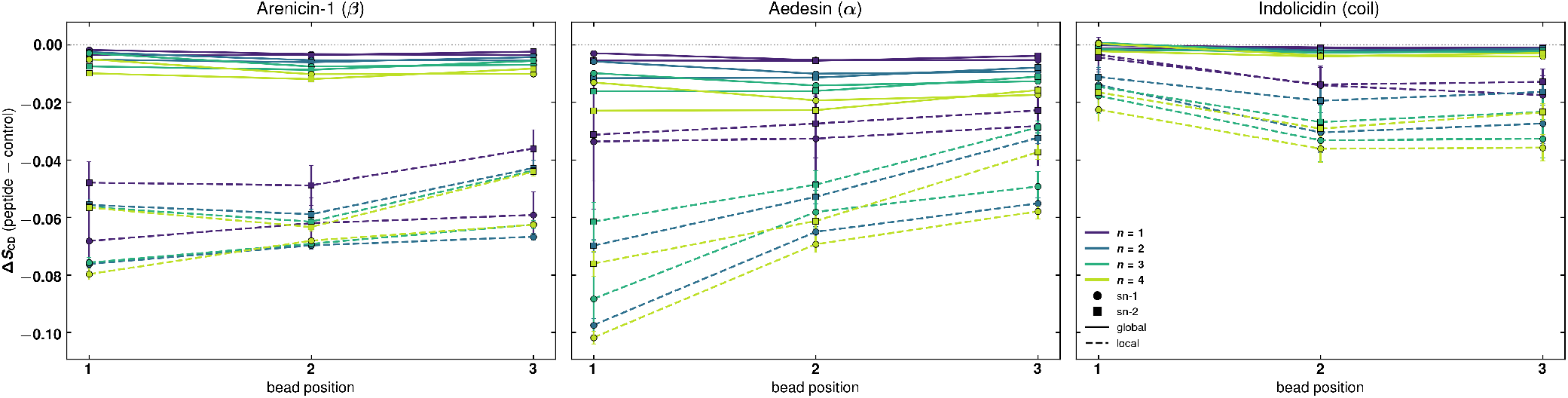
Change in acyl chain order parameter Δ*S*_CC_ (peptide minus control) on the peptide facing leaflet as a function of bead position along the sn-1 (circles) and sn-2 (squares) chains, for arenicin-1 (left), aedesin (center), and indolicidin (right). Colors denote peptide number n = 1 to n = 4. Solid lines show the global scope (all lipids in the leaflet) and dashed lines the local scope (lipids within 2.5 nm laterally of the nearest peptide center of mass). The global response scales with peptide length and grows with concentration, whereas the local per lipid response is largely set at n = 1, clusters for the two structured peptides independent of their length difference, and is markedly weaker for the disordered indolicidin. per peptide order parameter profiles for each chain are provided in Supplementary Figure S4.

The two scopes behave oppositely with respect to peptide length. The global Δ*S*_CC_, averaged over the peptide-facing leaflet and over all six tail bead positions at n = 4, was −0.019 for aedesin, −0.009 for arenicin-1, and −0.003 for indolicidin, an ordering that follows peptide length almost perfectly (Pearson *r* = −0.997 against residue count). This is the expected extensive signature: a longer peptide occupies a larger lateral footprint, recruits more lipids into that footprint, and therefore produces a larger bilayer-averaged chain disordering, independent of how strongly any individual lipid is perturbed. The global order parameter is thus, by construction, a length proxy and cannot by itself discriminate a structural-class mechanism from a size effect.

The local, per-lipid response does not follow length. The local Δ*S*_CC_ at n = 4, restricted to first-shell lipids, was −0.067 for aedesin and −0.062 for arenicin-1 but only −0.027 for indolicidin: the two structured peptides impose nearly identical per-lipid splay (a ratio of 0.93) despite a 15-residue difference in length, while the disordered peptide, intermediate in length, imposes less than half. Measured against a per-residue scaling, the deviations point in opposite directions—a diagnostic that no single monotonic length law can accommodate: arenicin-1 overperforms its length (effect ratio 0.93 versus length ratio 0.58 relative to aedesin) while indolicidin underperforms (effect ratio 0.44 versus length ratio 0.62 relative to arenicin-1). The local response therefore collapses onto the structured-versus-disordered axis rather than the length axis, mirroring at the single-lipid level the qualitative *K*_*A*_ dissociation of Section III B. The per-lipid splay is largest at the bead nearest the glycerol backbone and decays toward the chain terminus (for arenicin-1 at n = 4 the local sn-1 Δ*S*_CC_ values are −0.068, −0.057, and −0.063 from bead 1 to bead 3), the spatial signature of a perturbation seated at the top of the chains and pressing downward, as expected for a wedge acting from above the PO_4_ plane rather than a probe acting from within the bilayer core.

The concentration dependence reconciles the two scopes with the linear *K*_*c*_ softening. The per-lipid local response is largely established already at n = 1, retaining 86% of its n = 4 magnitude for arenicin-1, whereas the global response grows steadily with peptide number (for aedesin the magnitude of the global chain-averaged Δ*S*_CC_ increases from 0.005 at n = 1 to 0.019 at n = 4). The dose-dependent chain disordering therefore arises because additional peptides recruit additional lipids into perturbed footprints, not because each footprint lipid is splayed progressively harder. This is the chain-order counterpart of the concentration-invariant binding geometry established in Section III C and the cumulative-independent-perturbation interpretation of the linear *K*_*c*_ softening in Section III A.

Two conclusions follow that bound the claims made in the remainder of this work. First, the qualitative mode of chain coupling—expressed as whether a peptide imposes a strong per-lipid splay (structured) or a weak one (disordered)—is set by conformational state and is independent of peptide length, since the longest and intermediate-length peptides produce comparable per-lipid splay while the shortest-but-disordered peptide produces far less. Second, the present three-peptide design cannot separate the residual within-class magnitude difference between aedesin and arenicin-1 from their co-varying length, net charge, and wedge height; the two structured peptides differ by only 7% in per-lipid splay, and we accordingly do not interpret the alpha-versus-beta ordering as a clean structural-class magnitude law. For this reason, we report the bending and area compressibility moduli on a strict per-peptide basis (the dose-dependent slopes of Sections III A and III B) and against the geometric wedge descriptor, and we do not normalize the moduli by residue count, which would impose a per-residue scaling that the localization and local chain-order data jointly exclude.

### E. Direct measurement of bilayer thickness

The polymer brush analysis in Section III C predicted effective bilayer thickness reductions of 3.5%, 4.1%, and 8.9% for indolicidin, arenicin-1, and aedesin, respectively, at n = 4. To verify this prediction against a direct geometric observable, we computed two-dimensional thickness maps of the bilayer from the per-frame PO_4_– PO_4_ z-separation between leaflets, using a 4 Å × 4 Å lateral grid, with bin widths recomputed each frame to accommodate the NPT box fluctuations. Lipids were assigned to their original leaflet by an index-based block partitioning rather than a per-frame z-classification, ensuring that leaflet identity is preserved under any undulation or peptide-induced curvature. The analysis was performed over the same final 500 ns window used for the spectral and area-fluctuation analyses.

For the lipid-only control, the mean bilayer thickness was 3.983 ± 0.001 nm (mean ± SEM across five independent replicas), with a within-leaflet thickness variation of 0.030 nm representing the equilibrium undulation amplitude of the bilayer at 310 K. The polymer brush thickness inferred from the control K_*c*_ and K_*A*_ values reported in Sections III A and III B is 3.92 nm, in reasonable numerical agreement with the direct measurement and validating the use of the polymer brush relation as a semi-quantitative cross-check.

The bulk-averaged thickness varies weakly with concentration, and the magnitude of thinning at n = 4 orders as arenicin-1 (0.30%) > indolicidin (0.20%) > aedesin (0.15%), with mean thicknesses of 3.977 ± 0.001 nm for aedesin, 3.971± 0.001 nm for arenicin-1, and 3.975 ± 0.001 nm for indolicidin; the full concentration-resolved trend is shown in Figure S5. This ordering does not follow the K_*c*_ and K_*A*_ softening hierarchies. Rather than indicating a contradiction, this demonstrates that the bulk geometric thickness and the elastic moduli probe distinct physical aspects of the peptide-membrane interaction. The moduli are sensitive to the integrated stress response of the locally perturbed regions, dominated by chain splay and headgroup tilt induced by the supraphosphate wedge. The bulk geometric thickness, by contrast, is sensitive to direct displacement of PO_4_ headgroup beads by peptides that physically occupy the headgroup region, which scales inversely with the height of the peptide above the PO_4_ plane: aedesin, perched 0.49 nm above PO_4_, does not directly displace headgroups and produces the smallest bulk thinning despite the largest moduli effects; arenicin-1, partially embedded in the headgroup region at 0.22 nm above PO_4_, displaces a larger number of PO_4_ beads downward and produces the largest bulk thinning; indolicidin, sitting flush at PO_4_ with tryptophan anchors threading into the upper chain region, produces an intermediate bulk thinning through individual PO_4_ displacement at each anchor site. The three peptides therefore occupy distinct positions on a two-dimensional plane spanned by (i) wedge mechanical perturbation, which dominates the moduli, and (ii) direct headgroup displacement, which dominates the bulk geometric thickness.

The two-dimensional thickness maps (Figure 7) show that the bulk-averaged thickness is built from spatially heterogeneous local thinning. For aedesin at n = 3 and n = 4, discrete patches of thinning appear, with local thickness values as low as 3.85–3.90 nm compared to the unperturbed bulk of 3.98 nm. For arenicin-1, subtler thinning patches appear at higher concentration with smaller magnitude and reduced spatial extent. For indolicidin, the thickness maps remain essentially homogeneous with no visible thinning patches, consistent with a peptide that does not coherently displace lipids in the bilayer plane. The discrepancy between the polymer brush effective thinning (4–9%) and the bulk-averaged geometric thinning (<1%) reflects the spatial dilution of localized peptide perturbations over the ∼316 nm^2^ leaflet area. The moduli, being weighted by the locally perturbed chain region, recover the local thinning magnitude; the spatial mean, weighted equally over the entire leaflet, attenuates the local signal by a factor of order *A*_leaflet_*/A*_patch_.

**FIG. 7.**
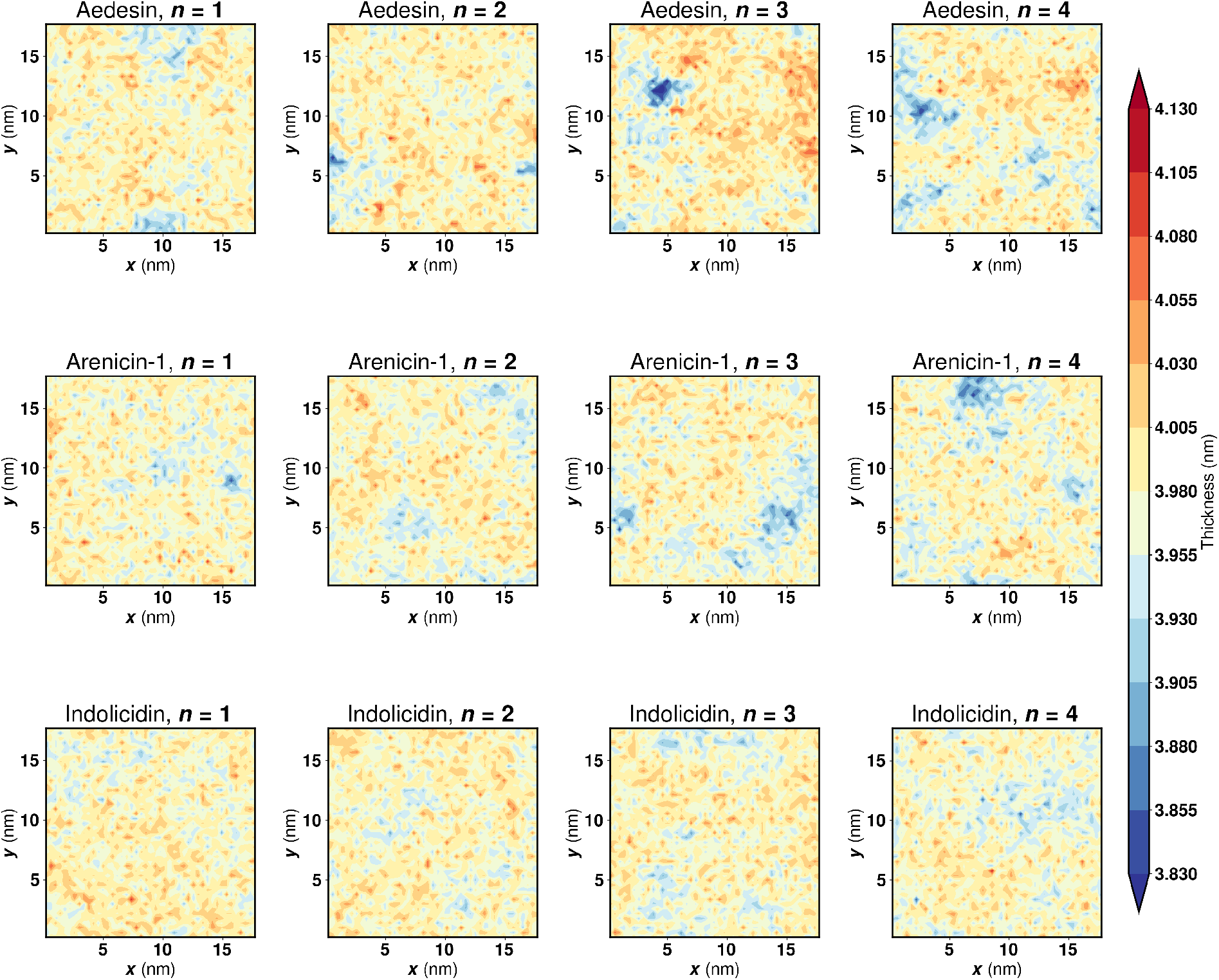
Two-dimensional bilayer thickness maps for the three AMP systems at n = 1 to n = 4. Maps show the local bilayer thickness averaged over the final 500 ns of each trajectory on a 4 Å × 4 Å grid. For aedesin at higher concentrations, discrete thinned patches appear at lateral positions corresponding to peptide-traffic-heavy regions integrated over the analysis window. For indolicidin, the maps remain essentially homogeneous, consistent with a peptide that does not coherently displace lipids in the bilayer plane.

The signature of local thinning that the bulk mean misses is preserved in the within-replica spatial standard deviation of the thickness, which quantifies the lateral heterogeneity of the bilayer surface. The SD grew systematically with peptide loading for all three peptides, from 0.031 nm at n = 1 to 0.032 nm at n = 4 for indolicidin (a 5% increase relative to its n = 1 value), from 0.032 to 0.036 nm for arenicin-1 (an 11% increase), and from 0.033 to 0.039 nm for aedesin (a 20% increase). This ordering of lateral-heterogeneity growth tracks the supraphosphate adsorption versus interfacial insertion hierarchy established in Section III C: aedesin, sitting highest above the PO_4_ plane with the largest lateral footprint, induces the strongest spatial inhomogeneity in bilayer thickness; arenicin-1 produces an intermediate effect; and indolicidin, inserting into the interface without coherently displacing lipids, produces the mildest heterogeneity growth. Notably, the SD increase for indolicidin is only marginally above the control SD of 0.030 nm, consistent with its inability to soften K_*A*_ reported in Section III B: a peptide that does not coherently displace lipids cannot generate substantial lateral thickness variation. The thickness SD therefore provides an independent, geometrically direct observable that distinguishes the supraphosphate adsorption and interfacial insertion binding modes, complementing the moduli-based and depth-based descriptors developed in the preceding sections.

### F. Spatial organization of peptides and lipids

The supraphosphate adsorption versus interfacial insertion hierarchy developed in Section III C predicts not only differential mechanical signatures but also differential spatial organization of peptides and lipids on the membrane surface. To test this prediction we computed two complementary radial distribution functions in the peptide-binding leaflet: a peptide-peptide RDF *g*_*pp*_(*r*) that probes peptide-peptide spatial correlations, and peptide-lipid RDFs *g*_*pL*_(*r*) for POPE and POPG separately that probe peptide-lipid spatial correlations. All distances were evaluated in the lateral xy plane under the minimum image convention; lipids were restricted to the peptide-binding (upper) leaflet via the same index-based assignment used for the thickness analysis. Both RDFs were normalized against the per-frame mean density of the partner species in the box, so *g*(*r*) = 1 corresponds to random spatial distribution.

The peptide-lipid RDFs (Figure 8) reveal a charge-correlated enrichment of POPG in the first solvation shell of each peptide. At n = 1, the first-peak amplitude of *g*_*p*,POPG_(*r*) at r ≈ 1.2 nm was 1.55 for aedesin, 1.38 for arenicin-1, and 1.22 for indolicidin, ordering precisely with the net peptide charge (+8, +6, +4 respectively). The corresponding *g*_*p*,POPE_(*r*) curves were depleted at small r for all three peptides, with the POPE deficit deepest and most spatially extended for aedesin, intermediate for arenicin-1, and shallowest for indolicidin. Both curves converged to unity beyond r ≈ 4 nm, indicating that the lipid-composition perturbation is local to the peptide first shell and the bulk POPE:POPG ratio is recovered at larger separations. The shape and amplitude of the first peak were essentially independent of peptide concentration, consistent with the saturated single-peptide binding mode established in Section III C. The POPG enrichment is electrostatic in origin and scales with peptide charge rather than with structural class: the bulk leaflet POPG density (153 lipids per 316 nm^2^) is sufficient that multiple peptides do not measurably compete for the available anionic lipid pool.

**FIG. 8.**
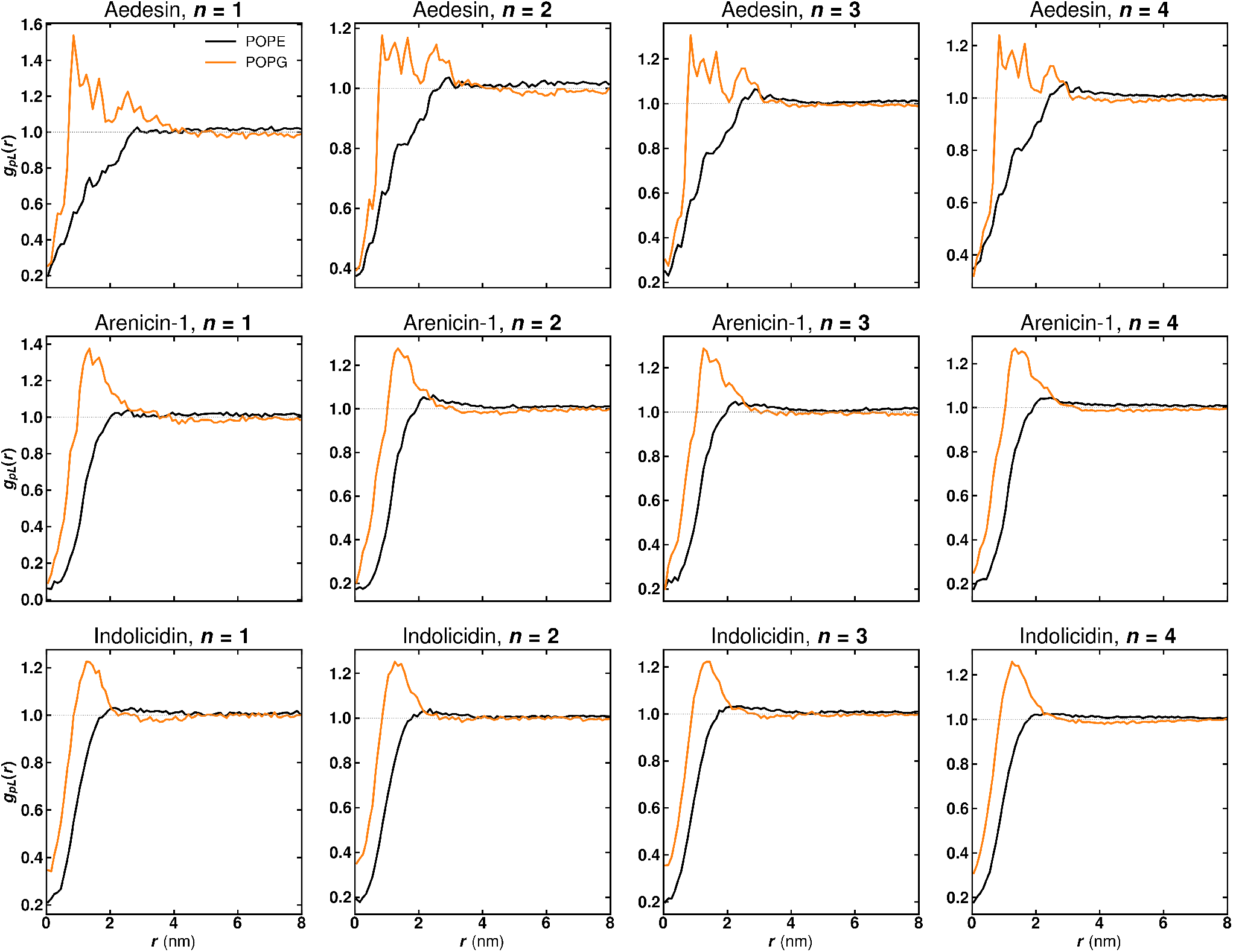
Peptide-lipid radial distribution functions *g*_*pL*_(*r*) for the three AMP systems at n = 1 to n = 4, computed separately for POPE (black) and POPG (orange) in the peptide-binding leaflet. The first-peak amplitude of *g*_*p*,POPG_(*r*) at r ≈ 1.2 nm orders with peptide charge: 1.55 for aedesin (+8), 1.38 for arenicin-1 (+6), and 1.22 for indolicidin (+4). The POPG enrichment is local to the peptide first shell and the bulk POPE:POPG ratio is recovered beyond r *≈* 4 nm. The dotted line marks *g*(*r*) = 1.

The peptide-peptide RDFs (Figure 9) reveal structural class specific differences in peptide-peptide spatial organization. At n = 2 and n = 3 all three peptides exhibit g_*pp*_(r) < 1 across the accessible distance range, indicating that peptide pairs predominantly occupy separations >8 nm and tend to avoid one another within the minimum-image window. At n = 4, however, aedesin separates from the other two peptides with a distinct short-range peak at r ≈1.5-2 nm reaching *g*_*pp*_(*r*) ≈1.5-1.8, suggestive of peptide crowding or a close-contact tendency between alpha-helical rods at high surface concentration; whether this signature reflects a kinetic encounter or a thermodynamic preference cannot be resolved on the simulation timescale considered here. arenicin-1 and indolicidin, in contrast, show g_*pp*_(r) ≲ 1 throughout the 0-3 nm range at n = 4, indicating no analogous short-range correlation. The aedesin short-range peak is consistent with the discrete thinning patches seen at n = 4 in the thickness maps (Figure 7): when peptides occupy nearby positions their individual wedge footprints overlap and produce localized thinned regions in the bilayer. The absence of an analogous signature for arenicin-1 and indolicidin is consistent with their compact and disordered geometries, respectively, neither of which presents the extended hydrophobic interface that would favor close-contact configurations at the membrane surface.

**FIG. 9.**
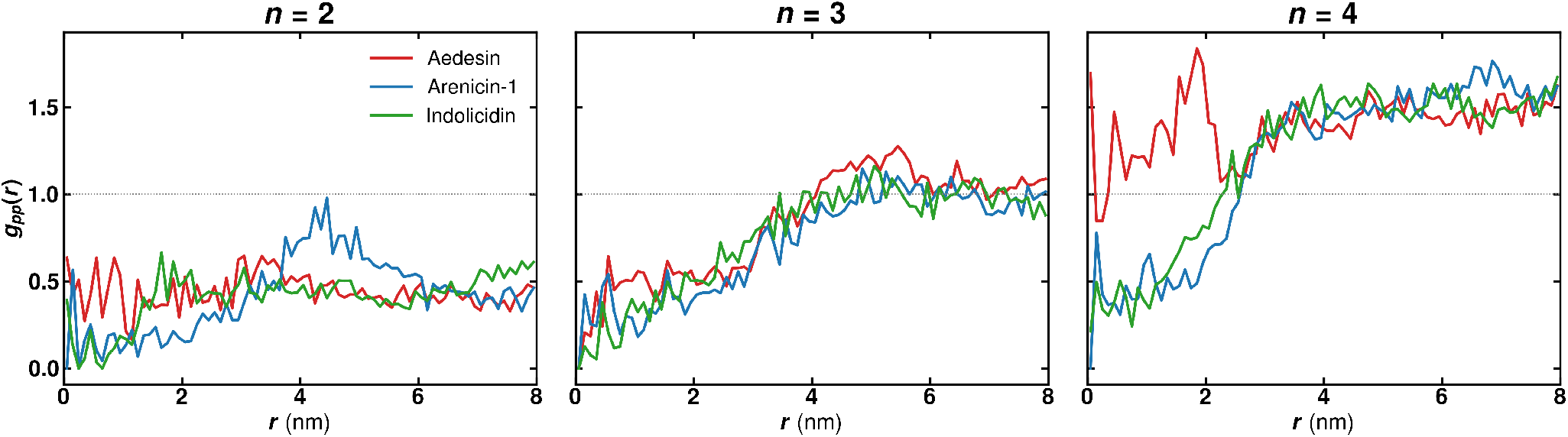
Peptide-peptide radial distribution functions *g*_*pp*_(*r*) for the three AMP systems at n = 2, 3, and 4. At n = 4, aedesin exhibits a short-range peak at r ≈ 1.5-2 nm reaching *g*_*pp*_(*r*) ≈ 1.5-1.8, suggestive of peptide crowding or a close-contact tendency at high surface concentration. Arenicin-1 and indolicidin show no analogous short-range correlation at any concentration. The dotted line marks *g*(*r*) = 1.

Taken together, the peptide-lipid and peptide-peptide RDFs provide two independent observables that corroborate the mechanistic framework developed in Sections III C and III E. The POPG enrichment ordering tracks peptide charge and identifies the electrostatic component of peptide-lipid association as charge-driven rather than structural class driven. The peptide-peptide short-range correlation, by contrast, is structural class specific, appearing only for the alpha-helical aedesin at the highest concentration studied and mirroring the localized thinned regions seen at n = 4 in the thickness maps.

### G. Dose dependent change in lipid packing

The mechanical, geometric, and spatial-organization analyses presented in Sections III A–III F characterize the bilayer response to peptide loading at the continuum and supramolecular levels. To complement this picture with a microstructural observable that probes lipid packing at the single headgroup scale, we computed leaflet resolved lipid packing defect distributions using the PackMem framework [20]. Packing defects are sites of imperfect lipid headgroup coverage that expose acyl chain material to solvent, and prior work has identified them as preferred anchoring sites for amphipathic membrane active peptides [21, 22]. The dose dependent evolution of the defect distributions therefore provides a microstructural counterpart to the bulk mechanical and geometric observables analyzed above.

For all three peptide systems, the upper (peptide facing) leaflet exhibited a progressive decrease in the number of deep defect sites with increasing peptide concentration, accompanied by a concomitant increase in the mean area of individual deep defects. In the arenicin-1 system, the mean deep defect count on the upper leaflet decreased from 141.7 in the control to 139.9, 137.9, 136.4, and 134.6 at concentrations of n = 1 through n = 4, respectively. Simultaneously, the mean individual deep defect area increased from 6.09 Å^2^ in the control to 6.57, 6.99, 7.31, and 7.68 Å^2^. The total deep defect area on the upper leaflet consequently increased from 863.7 to 919.0, 963.7, 997.3, and 1033.1 Å^2^, representing a cumulative increase of approximately 20% at the highest concentration.

The aedesin system exhibited analogous trends in defect remodeling, with the upper leaflet deep defect count decreasing from 141.7 to 140.1, 137.9, 136.0, and 134.3 at n = 1 through n = 4. The mean individual deep defect area rose from 6.09 to 6.32, 6.50, 6.83, and 7.03 Å^2^, and the total deep defect area increased from 863.7 to 886.1, 896.2, 928.9, and 944.1 Å^2^. While the absolute magnitude of defect area increase was somewhat smaller for aedesin than for arenicin-1 at comparable concentrations, the decrease in defect count proceeded at a similar rate for both peptides.

The indolicidin system showed qualitatively similar remodeling of the upper leaflet deep defects despite its disordered, surface-flush binding mode. The mean deep defect count on the upper leaflet decreased from 141.7 in the control to 140.9, 140.0, 138.8, and 137.9 at n = 1 through n = 4, while the mean individual deep defect area rose from 6.09 to 6.43, 6.83, 7.15, and 7.59 Å^2^. The corresponding total deep defect area on the upper leaflet increased from 863.7 to 906.0, 955.9, 992.0, and 1046.3 Å^2^, a cumulative increase of approximately 21% at the highest concentration that is comparable in magnitude to the arenicin-1 response.

This pattern, wherein fewer but individually larger deep defects coexist with an overall increase in total exposed area, indicates that peptides stabilize and expand existing deep defects rather than nucleating new ones. The physical interpretation is that AMPs, upon interacting with the membrane surface, anchor in the vicinity of pre-existing deep defects and through their hydrophobic components prevent the transient closure of these voids, effectively consolidating neighboring small defects into larger, more persistent ones [22]. It is pertinent to note here that the PackMem software cannot measure defect volume and there is a possibility that aedesin causes comparatively more softening than arenicin-1 because of larger defect volumes.

The per-frame distributions of defect counts and total deep defect areas (Figures S6 and S7) show systematic shifts in the upper leaflet histograms with peptide loading that are consistent with the mean-value trends reported above: the defect-count distributions shift modestly leftward and the total-area distributions shift cleanly to higher values, with the n = 4 distribution offset from control by an amount that exceeds the inter frame spread. The lower leaflet distributions remain essentially superimposed across conditions for all three peptides, confirming that the peptide induced perturbation is restricted to the peptide facing leaflet. The individual defect size distributions (Figure 10) reveal that the total-area shift is driven by an extension of the heavy tail of the deep defect *P* (*A*) on the upper leaflet, with peptide-loaded systems sustaining probability densities one to two orders of magnitude above control for defect areas in the 80-200 Å^2^ range. The lower leaflet deep defect distributions and the shallow defect distributions on the lower leaflet remain essentially invariant across conditions for all three peptides.

**FIG. 10.**
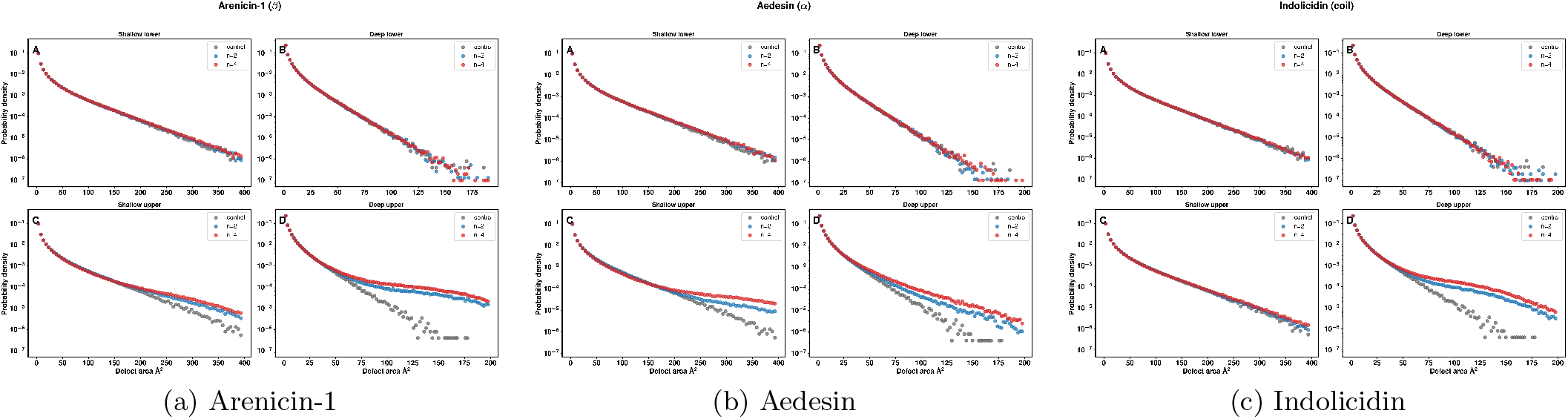
Probability density of individual lipid packing defect areas *P* (*A*) for the three AMP systems at the highlighted concentrations (control vs n = 2 vs n = 4), plotted on a log-y axis with fixed bin edges aligned to the PackMem 1 Å^2^ integer grid. Panel layout is as follows: (A) shallow lower, (B) deep lower, (C) shallow upper, (D) deep upper. The deep upper panels (D) show systematic extension of the heavy tail of *P* (*A*) with peptide loading for all three peptides, with peptide loaded systems sustaining probability densities one to two orders of magnitude above control in the 80-200 Å^2^ range. The shallow upper panels (C) show a similar tail extension for the structured peptides arenicin-1 and aedesin, but a markedly weaker shift for the disordered indolicidin whose shallow *P* (*A*) remains close to control across the full range. The lower leaflet panels (A, B) remain essentially overlapping across conditions for all three peptides, consistent with the leaflet-specific perturbation reported in the text.

The shallow defect profile on the upper leaflet, in contrast, reveals a structural class asymmetry that the deep channel alone does not. The two structured peptides exhibit the same consolidation signature on shallow defects as on deep, namely a decrease in shallow defect count combined with an increase in the mean individual shallow defect area, while the total shallow area remains essentially conserved. At n = 4 the upper leaflet shallow defect count dropped by 4.2% for arenicin-1 (Welch *p <* 10^−11^, Hedges’ *g* = −7.3) and 7.8% for aedesin (*p <* 10^−9^, *g* = −10.0), with concomitant increases in the mean individual shallow defect area of 3.9% for arenicin-1 (raw *p* = 2 × 10^−3^, *g* = +1.2) and 7.5% for aedesin (raw *p* = 10^−2^, *g* = +1.2); the corresponding shallow total areas changed by less than 1% in both cases and were not statistically resolved. The disordered indolicidin, by contrast, shows only a marginal 1.3% decrement in shallow defect count and no statistically resolvable change in the mean individual shallow defect area (+0.3%, *p* = 0.35, *g* = +0.3). The deep channel remodels for all three peptides with comparable magnitudes (total deep area increases of +19.6%, +9.3%, and +21.1% at n = 4 for arenicin-1, aedesin, and indolicidin respectively; all *p <* 10^−7^ after Bonferroni correction across 144 comparisons), so the asymmetry is specific to the shallow channel. The structural class signature in the defect response is therefore: structured peptides remodel both shallow and deep voids on the peptide facing leaflet, while the disordered peptide remodels only deep voids. This dichotomy is consistent with the per-bead z-distributions reported in Section III C: the structured peptides have a substantial fraction of their bulk residing in the headgroup region above the glycerol reference plane and therefore directly occlude shallow voids by mass occupancy, while indolicidin has its backbone at the PO_4_ plane and its sidechains predominantly below the glycerol level, leaving the intervening shallow defect probe depth only weakly perturbed.

The leaflet resolved defect remodeling closes the multi scale picture developed across Sections III A–III F: peptides act exclusively at the binding leaflet, where they consolidate preexisting defects into a smaller number of larger features through their surface bound footprints. The deep defect consolidation is shared across all three structural classes and reflects a generic interfacial anchoring behavior, but the shallow defect response discriminates structured from disordered AMPs and provides an independent microstructural readout of the supraphosphate adsorption versus interfacial insertion dichotomy established in Section III C. The mechanical softening, geometric thinning, and lateral spatial organization (Sections III B–III F), together with the shallow defect response, therefore discriminate the three structural classes through their differential supraphosphate occupancy and lateral footprint. The defect distributions complement rather than recapitulate the moduli based and geometric descriptors developed earlier in this work.

## IV. DISCUSSION AND CONCLUSION

This work establishes a quantitative, dose-dependent picture of how three structurally distinct antimicrobial peptides perturb the mechanical and microstructural properties of a model Gram-negative bacterial inner membrane. All three peptides soften the bending modulus K_*c*_ monotonically with surface concentration, but the per-peptide softening rate spans a threefold range that orders systematically with peptide structural class: −1.39 ± 0.09 k_B_T per peptide for the alpha-helical aedesin, −0.66 ± 0.04 k_B_T per peptide for the beta-hairpin arenicin-1, and −0.44 ± 0.01 k_B_T per peptide for the disordered indolicidin. The area compressibility modulus K_*A*_ dissociates qualitatively from this response for indolicidin alone, softening for the two structured peptides while remaining statistically indistinguishable from the control for the disordered one. Independent geometric, spatial-organization, and microstructural observables corroborate the same underlying picture from complementary directions, and together they recast the mechanical hierarchy as the macroscopic output of two distinct membrane binding modes.

The organizing principle that emerges from the depth analysis is a supraphosphate adsorption versus interfacial insertion dichotomy. Aedesin and arenicin-1, with their secondary structure enforced by the ElNeDyn elastic network, present rigid amphipathic surfaces to the membrane and sit measurably above the PO_4_ plane (0.49 nm and 0.22 nm, respectively), acting as supraphosphate wedges whose rigid volumes force the underlying acyl chains to splay outward and produce both K_*c*_ and K_*A*_ softening. Indolicidin, by contrast, sits flush with PO_4_ and threads its tryptophan side chains into the interstitial space between lipid headgroups without coherently displacing them, perturbing local lipid tilt and curvature stress sufficiently to soften K_*c*_ but not strongly enough to disrupt the integrated chain packing at the bilayer core. The depth ordering of the three peptides is the inverse of what a naive insertion-depth picture would predict, and the same inversion is recovered in the bulk geometric thickness measurement: aedesin produces the smallest bulk thinning despite the largest moduli effects, while arenicin-1 produces the largest bulk thinning by virtue of its partial embedment in the headgroup region. This is not a contradiction but a confirmation that bulk thickness and elastic moduli probe distinct physical aspects of the perturbation, with the former tracking direct PO_4_ displacement and the latter tracking the integrated chain splay induced by the wedge.

The per-bead z-distributions in Figure 4 make this geometric distinction quantitative and provide a finer-grained view than the center-of-mass coordinate alone. At n = 4, aedesin retains 85% of its beads above the PO_4_ plane, arenicin-1 retains 67%, and indolicidin keeps only 44% of its beads above PO_4_. The disordered peptide therefore samples below-PO_4_ space with more than half of its beads, dominated by deep penetration of tryptophan and arginine side chains, even as its backbone remains essentially flush with PO_4_. This decoupling between backbone and side-chain occupancy is the geometric signature of conformational disorder at the interface and underpins the entire framework: the structured peptides occupy headgroup-region volume coherently with both backbone and side-chain mass, while indolicidin distributes its mass discontinuously across the interface with the backbone above and the side chains threading through and below.

The dissociation of K_*c*_ and K_*A*_ for indolicidin is, to our knowledge, a previously unreported mechanical signature that discriminates these two binding modes more cleanly than either modulus alone. The two elastic moduli, though both global descriptors of the bilayer, sample different regions of the bilayer cross-section: K_*c*_ integrates contributions from the entire bilayer thickness with substantial weight on peripheral lipid tilt and curvature stress, whereas K_*A*_ is dominated by entropic chain packing at the core. A peptide that perturbs interfacial mechanics without coherently displacing lipids in the membrane plane will therefore soften K_*c*_ but leave K_*A*_ largely unaffected, which is precisely what we observe for indolicidin. The acyl chain order parameter analysis (Section III D) supplies the single-lipid corroboration of this distinction: the structured peptides impose a strong per-lipid chain splay that the disordered peptide does not, and because this per-lipid measure separates the disordered peptide from the structured ones independently of peptide length, it establishes that the qualitative coupling mode is set by conformational state rather than by molecular size. The polymer brush relation 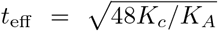 collapses the two moduli into a single effective thickness descriptor that recovers the structural-class ordering quantitatively, with the indolicidin effective thinning of 3.5% at n = 4 generated almost entirely by the K_*c*_ reduction and no contribution from K_*A*_. We suggest that K_*A*_ be included as a standard observable in computational studies of AMP-membrane interactions whenever the peptide is suspected to be conformationally disordered or weakly amphipathic.

The strong pooled correlation between per-replica K_*c*_ and the corresponding replica-averaged peptide position above PO_4_ (Pearson r = − 0.84, p < 10^−30^, N = 116) provides direct quantitative support for the geometric framework. Within each structural class the correlation is substantially weaker, with position varying by less than 0.2 nm and per-replica K_*c*_ varying by 4–5 k_B_T, indicating that the height above PO_4_ is a class-level rather than a continuous predictor of softening. The relevant physical descriptor is the structural-class identity, encoded geometrically by the volume and lateral footprint of the peptide body residing in the headgroup-water region, with within-class variations dominated by configurational sampling at fixed n. The growth of inter-replica K_*c*_ variance with peptide number reflects the increasing diversity of accessible metastable peptide-membrane configurations at higher loading, sampled stochastically across independent replicas, rather than any change in the underlying perturbation mechanism.

The K_*c*_ softening is linear in peptide number over the range probed here, with statistically significant slopes (p ≤ 0.004) for all three peptides and no evidence of saturation or inflection. Combined with the depth analysis showing that mean peptide binding geometry is concentration-invariant, this linearity supports a model in which surface-bound AMPs act as independent mechanical perturbations whose individual contributions add cumulatively up to the highest concentration probed. There is no evidence for cooperative insertion, for a critical surface density at which the perturbation mechanism changes character, or for any concentration-dependent reorientation of bound peptides. Whether the linear dose response extends to higher concentrations remains an open question: saturation of available binding sites, the onset of cooperative peptide-peptide interactions, or transitions to insertion or pore-forming regimes are all plausible at concentrations beyond the present study.

The peptide-peptide radial distribution functions reveal a structural-class-specific lateral organization that emerges only at the highest concentration. At n = 2 and n = 3 all three peptides exhibit *g*_*pp*_(*r*) < 1 throughout the accessible distance range, indicating mutual avoidance. At n = 4, however, aedesin develops a short-range peak at r ≈ 1.5–2 nm reaching *g*_*pp*_(*r*) ≈ 1.5– 1.8, suggestive of a close-contact tendency between alpha-helical peptides, while arenicin-1 and indolicidin show no analogous signature. The aedesin close-contact peak coincides geometrically with the discrete thinning patches resolved in the two-dimensional thickness maps at the same concentration, consistent with overlapping wedge footprints producing locally thinned bilayer regions. Whether this peak reflects a thermodynamic preference for close-contact configurations or a kinetic encounter that has not yet relaxed on the simulation timescale cannot be resolved here. The class specificity of this peptide-peptide correlation marks the onset of a potentially cooperative regime accessible only to the extended alpha-helical structural class and motivates longer simulations at higher concentrations.

The leaflet-resolved lipid packing defect analysis provides the microstructural counterpart to the bulk observables and reveals a two-channel signature that further discriminates the structural classes. The deep-defect channel on the peptide-facing leaflet remodels universally across all three peptides: defect count decreases, mean individual defect area increases, and the heavy tail of P(A) extends to substantially larger area values, producing cumulative total deep-defect area increases of 19.6% for arenicin-1, 9.3% for aedesin, and 21.1% for indolicidin at n = 4. This consolidation of pre-existing deep defects into a smaller number of larger, longer-lived features reflects a generic surface-anchoring behavior in which the hydrophobic components of any surface-bound AMP, regardless of structural class, prevent the transient closure of preexisting voids in their immediate neighborhood. The shallow-defect channel, in contrast, distinguishes structured from disordered peptides cleanly: aedesin and arenicin-1 generate the same consolidation signature on shallow defects as on deep ones, with significant decreases in shallow defect count (7.8% and 4.2% at n = 4, respectively) and increases in the mean individual shallow defect area (7.5% and 3.9%), while indolicidin produces only a marginal 1.3% decrement in shallow count and no statistically resolvable change in mean shallow defect area. This shallow-channel asymmetry is the microstructural counterpart of the K_*A*_ dissociation and finds its mechanistic origin in the per-bead z-distributions: the structured peptides have a substantial fraction of their bulk residing in the headgroup region above the glycerol reference plane and therefore directly occlude shallow voids by mass occupancy, while indolicidin has its backbone at the PO_4_ plane and its side chains predominantly below the glycerol level, leaving the intervening shallow-defect probe depth only weakly perturbed. The structural-class signature in the defect response is therefore: structured peptides remodel both shallow and deep voids on the peptide-facing leaflet, while the disordered peptide remodels only deep voids. The caveat that PackMem cannot resolve defect volume, only the projected two-dimensional area, should be noted; the systematically larger K_*c*_ softening of aedesin relative to arenicin-1 at comparable defect-area perturbations may partially reflect this hidden volume dimension.

The defect remodeling results connect to a broader question in membrane biophysics: whether membrane-active amphipathic molecules sense the geometric curvature of a membrane or the lipid packing defects that are enhanced at curved surfaces [23, 25, 28]. Prior work has established that packing defects, internal membrane stresses, and elastic moduli are not independent quantities but different manifestations of the same underlying lateral pressure profile [9]. The relationship between curvature and defects has been explored extensively in one direction: how membrane curvature creates packing defects that recruit amphipathic proteins. The present results address the reverse direction by showing that AMPs remodel packing defects in ways that correlate with, and may mechanistically drive, changes in the bending modulus. The deep-defect consolidation observed for all three structural classes increases the total area of exposed hydrophobic surface on the peptide-facing leaflet, which is equivalent to a local reduction in the lateral pressure at the chain-headgroup interface; such a reduction, by altering the second moment of the lateral pressure profile, would directly reduce K_*c*_. The observation that aedesin produces the largest K_*c*_ softening despite a smaller total defect-area increase than indolicidin points to defect depth or volume, rather than projected area alone, as the mechanically relevant variable, consistent with the suggestion by Campelo and Kozlov [28] that the full depth-resolved stress profile, not just interfacial defects, governs the elastic response. Establishing whether the defect–K_*c*_ correlation is causal or reflects a common upstream origin in the lateral pressure redistribution will require direct measurement of the leaflet-resolved lateral pressure profile as a function of peptide dose, an analysis that the present simulation framework can support in future work.

Taken together, the structural-class hierarchy of mechanical softening rates, the K_*c*_–K_*A*_ decoupling for the disordered peptide, and the shallow-channel defect asymmetry suggest a design principle for engineered AMPs. Conformational rigidity, by stabilizing a fixed amphipathic geometry at the membrane surface, prevents the time averaging over a broad conformational ensemble that attenuates the wedge effect for disordered peptides. A peptide that is structurally constrained to maintain a defined amphipathic helical fold should therefore produce a stronger and more reproducible supraphosphate wedge perturbation than its unconstrained counterpart on a per-peptide basis. Conformational rigidity alone is not the whole story: the two structured peptides differ in their per-peptide softening, with the extended aedesin helix perturbing the bilayer more than the compact arenicin-1 hairpin. We are cautious about attributing this residual difference to the geometric class of the fold alone, because across these two peptides the fold geometry covaries with peptide length, net charge, and supraphosphate wedge height, and a three-peptide design cannot separate them; the acyl chain order parameter analysis shows that the per-lipid splay imposed by the two structured peptides differs by only 7%, so the larger bulk effect of aedesin reflects chiefly its larger footprint rather than a stronger per-lipid perturbation. The design principle we therefore advance is the conservative one: conformational restriction maximizes the per-peptide supraphosphate wedge perturbation relative to a disordered counterpart of similar composition. Stapled peptides, in which a hydrocarbon crosslink locks the alpha-helical conformation, offer a direct experimental realization of this principle and, by holding sequence and length fixed while varying only conformational rigidity, would also isolate the rigidity contribution from the length and charge covariates that the present comparison cannot disentangle. Experimental tests through vesicle-based flicker spectroscopy or micropipette aspiration on stapled and unstapled AMP analogs would constitute a direct validation of this prediction.

Several caveats are appropriate. The ElNeDyn elastic network restraints employed for aedesin and arenicin-1 preserve their native secondary structure and prevent the partial unfolding and chain insertion events observed for aedesin in atomistic simulations [21]. The present results therefore characterize the surface-adsorbed regime relevant to the early stages of AMP-membrane interaction, prior to any insertion or pore-forming events. The ElNeDyn constraint is, however, a reasonable computational model for synthetically constrained peptides such as stapled or cyclized analogs, lending the design inference above a defensible interpretation. The microsecond timescale of the present simulations may be insufficient to fully sample the configurational space at n = 4, as suggested by the inter-replica K_*c*_ variance growth at higher loading; this is partially mitigated by the use of multiple independent replicas per condition. The model membrane (binary POPE:POPG 70:30) captures the canonical inner-leaflet lipid composition of Gram-negative bacteria but omits lipopolysaccharide, membrane proteins, and the lipid asymmetry of the real bacterial envelope; whether the structural-class hierarchy of mechanical softening rates persists in more compositionally complex membranes remains to be tested. The three peptides differ simultaneously in conformational state, chain length, and net charge, and a three-peptide panel cannot statistically regress these covariates against one another; we have therefore anchored the mechanistic claims on observables that dissociate from length and charge (the qualitative K_*A*_ dissociation, the inverted insertion-depth ordering, and the per-lipid chain-order response, all of which separate the disordered peptide from the structured ones independently of length) and have explicitly declined to interpret the residual aedesin-versus-arenicin-1 magnitude difference as a pure structural-class effect. The stapled-versus-unstapled analog experiment proposed above is the design that removes the length and charge covariates by construction. Finally, the spectral elastic moduli reported here have been validated through the polymer brush consistency relation but have not been benchmarked against independent experimental measurements on the same lipid composition with the same peptides; experimental validation through flicker spectroscopy or micropipette aspiration on POPE:POPG vesicles doped with the three peptides studied here would substantially strengthen the present conclusions.

In summary, the bending modulus softens monotonically with peptide loading for all three AMPs at rates that order alpha-helical > beta-hairpin > disordered and span a threefold range across structural classes, while the area compressibility modulus dissociates qualitatively from the bending response for the disordered peptide alone. The supraphosphate adsorption versus interfacial insertion framework that emerges from the depth and per-bead z-distribution analyses organizes the entire dataset: structured peptides act as supraphosphate wedges that coherently displace lipid material in the headgroup region and soften both moduli, while the disordered peptide threads into the interface without coherent displacement and softens K_*c*_ alone. Independent observables drawn from bulk thickness analysis, lateral heterogeneity growth, peptide-peptide spatial organization, and the two-channel defect remodeling signature corroborate this framework from complementary directions, with the shallow versus deep defect asymmetry providing a particularly clean microstructural readout that mirrors the K_*A*_ dichotomy. The structural-class hierarchy supports a design principle in which conformational restriction maximizes the supraphosphate wedge perturbation per peptide relative to a disordered counterpart, motivating experimental tests on stapled-peptide AMP analogs that would hold sequence and length fixed while varying only conformational rigidity. Together, these findings establish a multi-scale, mechanistically grounded framework con-necting AMP structural identity, surface concentration, and the resulting perturbation of membrane mechanical and interfacial properties.

## V. ACKNOWLEDGMENTS

All simulations in this work have been carried out on the Kamet HPC cluster at The Institute of Mathematical Sciences, Chennai, India.

## VI. SUPPORTING INFORMATION

## Supplementary Information for

### S0.1. Lateral pressure profile analysis

#### 1. Method

Lateral pressure profiles were computed using a custom Python implementation of the Harasima convention [1], since native support for spatially resolved stress is unavailable in modern GROMACS and the GROMACS-LS extension [2] is incompatible with MARTINI 3 topologies. For planar bilayers, the Harasima and Irving-Kirkwood [3] conventions yield identical values for the integrated surface tension and for the moments of the stress profile [4]. The Harasima convention was selected for its simpler implementation, since lateral force contributions are assigned in equal halves to the slabs containing each interacting particle, avoiding the contour integration required by Irving-Kirkwood.

The implementation evaluates the nonbonded configurational stress arising from MARTINI 3 Lennard-Jones and reaction-field Coulomb interactions. Type-pair sigma and epsilon parameters were extracted from the [nonbond_params] section of the MARTINI 3 main itp file, since MARTINI 3 stores all interaction parameters there rather than in [atomtypes]. Pair lists within the cutoff were generated per frame using MDAnalysis [5, 6], with intramolecular exclusions constructed from the bond topology. The pairwise force evaluation and slab assignment were implemented in a numba [7] compiled kernel for performance.

Bonded forces (bonds, angles, dihedrals) and the kinetic stress contribution were omitted from the implementation. The kinetic contribution is isotropic and cancels in the lateral stress asymmetry. The bonded contribution is invariant across conditions at fixed lipid composition, so the concentration dependent difference profiles between peptide loaded and control systems are dominated by the nonbonded contributions; however, the bonded contribution is not negligible in absolute terms in the headgroup region, where it partially cancels the nonbonded torque. The present implementation therefore provides a quantitatively reliable view of the *shape* and *leaflet localization* of the concentration dependent stress perturbation, but cannot be used to extract absolute spontaneous curvature values. Such a quantitative extraction would require the full local stress tensor with bonded contributions, accessible through a CHARMM36 atomistic reference simulation analyzed with GROMACS-LS [2]; that benchmark is beyond the scope of the present work.

To suppress the broadening of profile features caused by thermal undulations of the bilayer in the lab frame, each frame was recentered on the bilayer midplane using the circular mean of the PO_4_ bead z-coordinates. For the box size and bending modulus regime relevant here ( ∼ 18 nm box, *K*_*c*_ ∼ 10–16 *k*_*B*_*T*), the residual undulation-induced error in the time-averaged Harasima profile is negligible relative to statistical noise. The lateral stress at slab z was computed as *s*(*z*) = *P*_*L*_(*z*) − *P*_*N*_, where the lateral component *P*_*L*_(*z*) is computed via the Harasima decomposition and the normal component *P*_*N*_ is taken as the constant applied pressure (1 bar) for the tensionless bilayer. Per-replica outputs include the slab z-coordinates, the lateral stress profile, and the integrated surface tension as a validation check, which lies near zero for all tensionless bilayer conditions. The analysis was performed over the same final 500 ns window used for the spectral and area-fluctuation analyses.

#### 2. Lateral pressure profiles

The lipid only control exhibits the canonical bilayer stress profile (Figure S1, dashed lines): positive lateral pressure peaks of ∼1000 bar at the two headgroup regions (*z* = ± 2 nm), flanking a negative trough at the acyl-chain interface that arises from the surface tension at the hydrophobic-hydrophilic boundary. The profile is symmetric about the midplane after time averaging and integrates to a surface tension of ∼ 0 bar nm, confirming the tensionless nature of the simulated bilayer.

In the peptide loaded systems, the perturbation to the lateral stress profile is strictly confined to the peptide facing leaflet (the upper leaflet, *z >* 0, by convention), with the distal leaflet remaining essentially superposed on the control profile across all concentrations for all three peptides (Figure S1). This one-sided character of the perturbation directly corroborates the leaflet-specific remodeling reported for the bilayer thickness maps (Section III D of the main text), the upper leaflet packing defect distributions (Section III F), and the peptide-lipid radial distribution functions (Section III E).

The peptide-induced difference profiles Δ*s*(*z*) = *s*_peptide_(*z*) − *s*_control_(*z*), shown in Figure S2, isolate the concentration dependent component of the stress redistribution and reveal a structural class dependent localization that matches the supraphosphate adsorption versus interfacial insertion framework developed in Section III C of the main text. For 1G89 (indolicidin, disordered), Δ*s*(*z*) develops a pronounced negative dip centered at *z* ≈ +1.5 nm, slightly inboard of the PO_4_ headgroup peak, with a magnitude that grows monotonically from ∼ −1000 bar at n = 1 to ∼ −4000 bar at n = 4. The inboard location of the dip is consistent with the per-bead z-distributions in Figure 3 of the main text, which place the tryptophan and arginine side chains of 1G89 substantially below the PO_4_ plane. For 2JSB (arenicin-1, beta-hairpin), Δ*s*(*z*) shows a narrower and shallower dip localized at the PO_4_ headgroup plane itself (*z* ≈ +2 nm), reaching ∼ − 1500 bar at n = 4. The location and width of the 2JSB feature are consistent with the partial embedment of the compact beta-hairpin within the head-group region (mean position 0.22 nm above PO_4_). For 2MMM (aedesin, alpha-helical), the difference profile remains close to zero across all concentrations, with only small (∼ ± 100 bar) fluctuations distributed across the upper leaflet. This is consistent with the 2MMM center of mass sitting 0.49 nm above the PO_4_ plane, in the headgroup-water region, where the elastic network constrained alpha-helical rod displaces water rather than perturbing the chain packing stress directly.

The ordering of the depth and amplitude of the Δ*s*(*z*) dip therefore recovers the same structural class hierarchy that organizes the bulk mechanical and microstructural observables in the main text: 1G89 sits deepest and produces the largest stress redistribution at the interfacial-insertion depth; 2JSB sits at the headgroup plane and produces an intermediate redistribution localized to the headgroup slab; 2MMM sits above the headgroups and produces no resolvable redistribution within the non-bonded stress profile.

#### 3. Caveats

Three caveats merit explicit statement. First, as noted above, the absolute magnitudes of Δ*s*(*z*) and any moments derived from it are not quantitatively comparable to spontaneous curvature values extracted from full-stress implementations. The first moment ∫*z* Δ*s*(*z*) *dz* computed from the present profiles overstates the true monolayer torque density by the bonded compensation factor, which is not estimated here. The shape and concentration scaling of Δ*s*(*z*) remain physically meaningful, but the integrated moments do not. Second, the 1G89 system exhibits a sign change in the integrated first moment between n = 2 and n = 3, while the Δ*s*(*z*) dip itself remains monotonically deepening. This sign change reflects a redistribution of stress between the supra-PO_4_ and sub-PO_4_ slabs at intermediate loading, possibly associated with a density-driven reorganization of the tryptophan-arginine side chain pattern, but the precise mechanism cannot be resolved from the present profiles alone. Third, the leaflet localization of the perturbation reported here is robust across all replicas after correcting for the per-replica orientation of the peptide-bound leaflet relative to the simulation box; this orientation correction is purely a post-processing step and does not affect the shape or amplitude of the leaflet-resolved Δ*s*(*z*) traces.

In aggregate, the lateral pressure profile analysis provides an independent z-resolved observable that corroborates the supraphosphate adsorption versus interfacial insertion framework established in the main text. The shape and depth of the perturbation order the three structural classes consistently with all other observables, while the strict one-sided character of the stress redistribution provides direct mechanical confirmation that the peptide induced perturbation is confined to the binding leaflet.

---

**FIG. S1.**
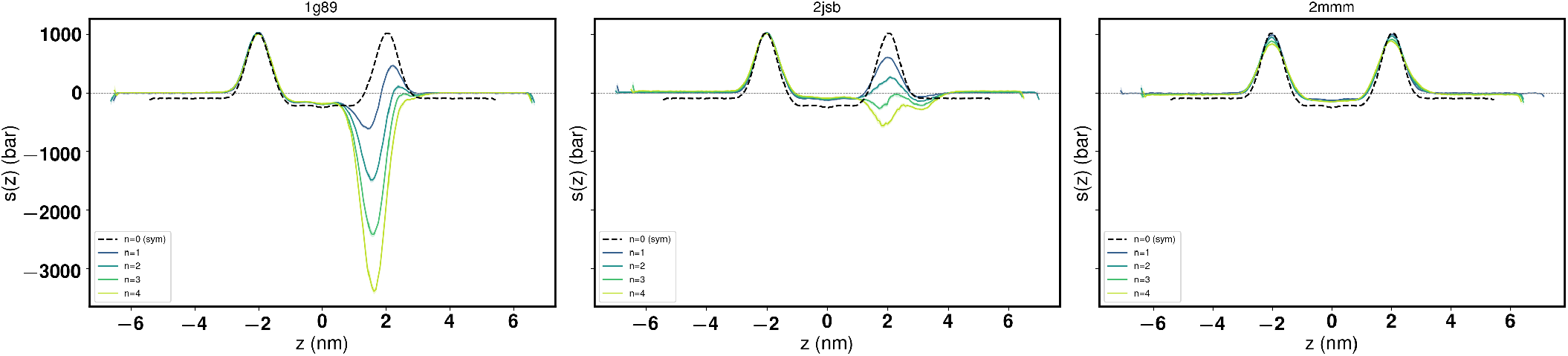
Time-averaged lateral pressure profiles *s*(*z*) for the three AMP systems at n = 1 to n = 4 peptides per leaflet. Solid colored lines are mean profiles across replicas (shaded bands: standard error of the mean); the dashed black line in each panel is the symmetrized lipid only control. The peptide binding leaflet is taken at *z >* 0 by convention; the distal leaflet at *z <* 0 remains essentially superposed on the control for all peptides and concentrations, demonstrating that the peptide induced stress perturbation is confined to the binding leaflet.

**FIG. S2.**
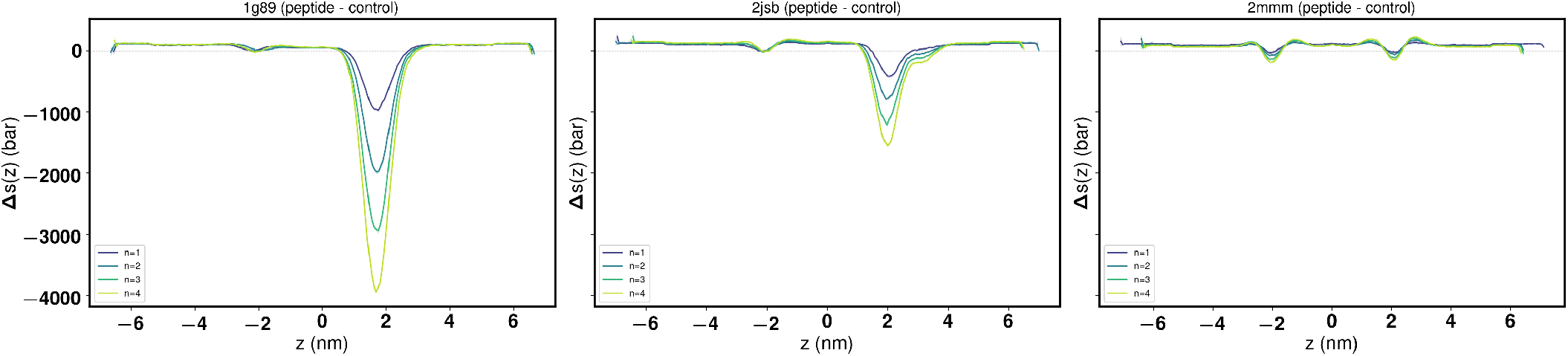
Peptide-induced difference profiles Δ*s*(*z*) = *s*_peptide_(*z*) − *s*_control_(*z*) for the three AMP systems at n = 1 to n = 4. The structural class ordering recovers the supraphosphate adsorption versus interfacial insertion hierarchy of the main text: 1G89 (disordered) produces the deepest perturbation, localized inboard of the PO_4_ plane consistent with side-chain insertion into the acyl chain region; 2JSB (beta-hairpin) produces an intermediate perturbation centered at the PO_4_ plane consistent with partial headgroup embedment; 2MMM (alpha-helical) produces no resolvable perturbation within the nonbonded stress profile, consistent with its position 0.49 nm above the PO_4_ plane in the headgroup-water region. The distal leaflet (*z <* 0) shows no peptide-induced perturbation for any system.

**FIG. S3.**
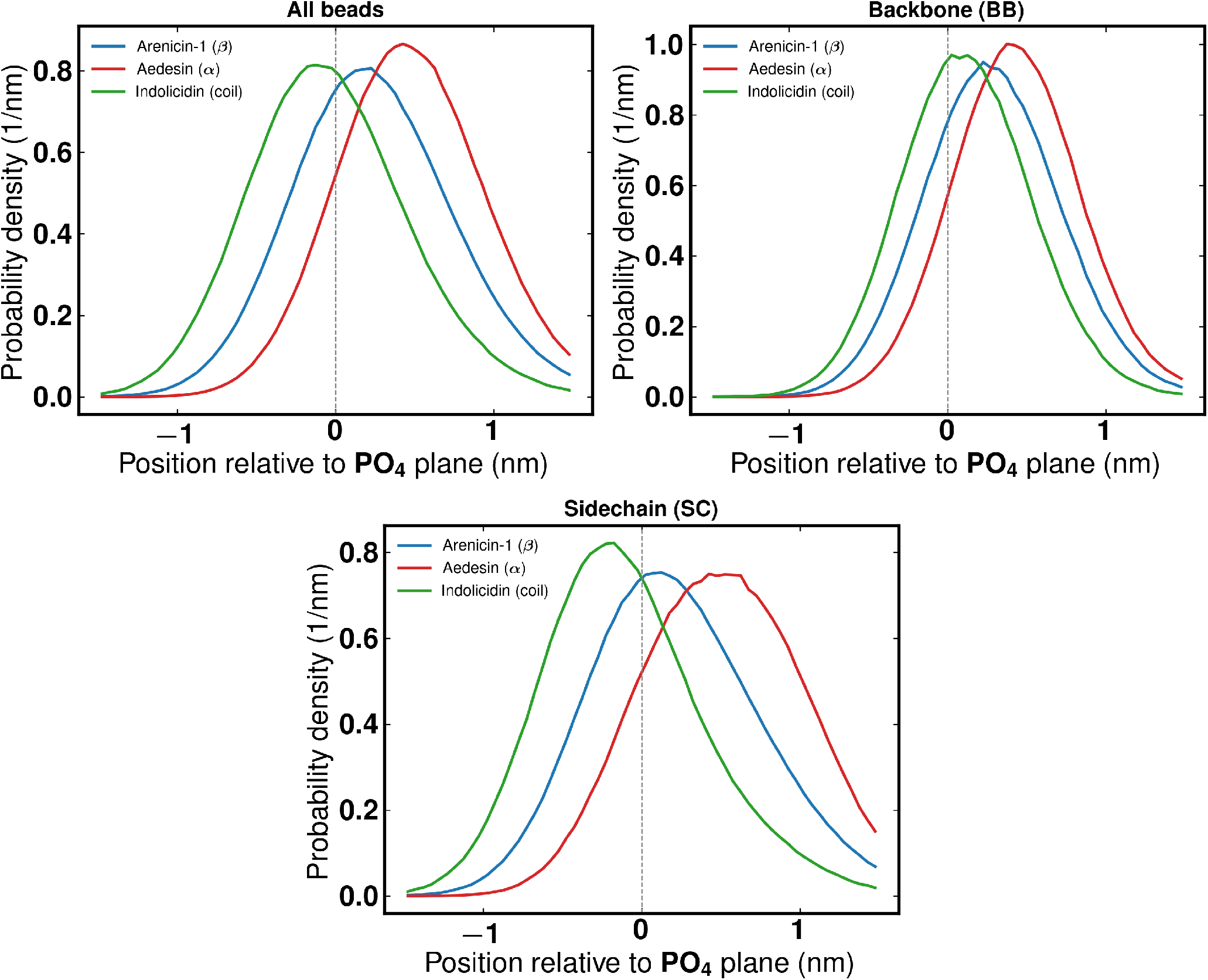
Per-bead position distributions relative to the peptide facing PO_4_ plane at n = 4 peptides per leaflet, for arenicin-1 (blue), aedesin (red), and idolicidin (green). The x-axis reports signed position in nm, with positive values indicating residence above PO_4_ (headgroup-water region) and negative values indicating residence below PO_4_ (toward the acyl chain region); the dashed vertical line marks the upper leaflet PO_4_ reference plane. Left panel shows all coarse-grained beads pooled; center panel shows backbone beads (BB) only; right panel shows sidechain beads (SC) only. Distributions are normalized to unit area. Aedesin and arenicin-1 keep the large majority of their beads above the PO_4_ plane (85% and 67% respectively), consistent with their supraphosphate adsorption mode. Indolicidin has 56% of its beads below PO_4_, with the sidechain distribution (SC fraction below PO_4_ : 62%) extending substantially further into the membrane than the backbone (BB fraction: 39%), reflecting the interfacial insertion of its tryptophan and arginine residues while the backbone remains flush with the headgroup plane.

**FIG. S4.**
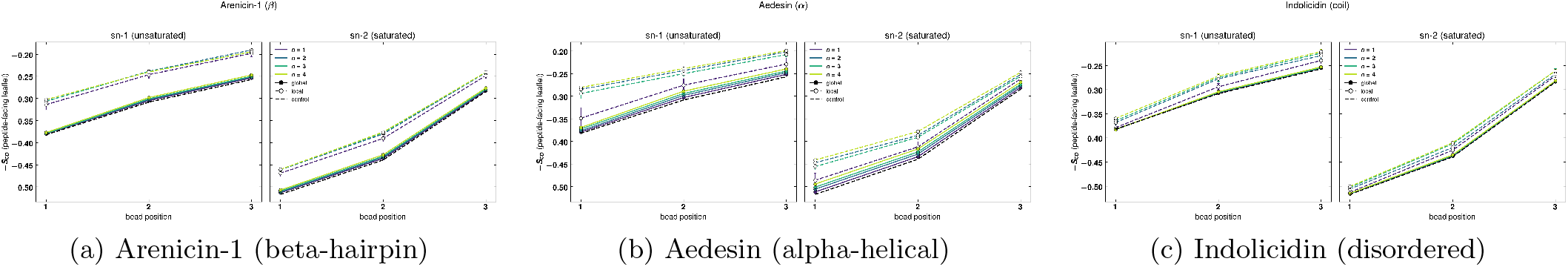
Per-chain SCD for each peptide class.

**FIG. S5.**
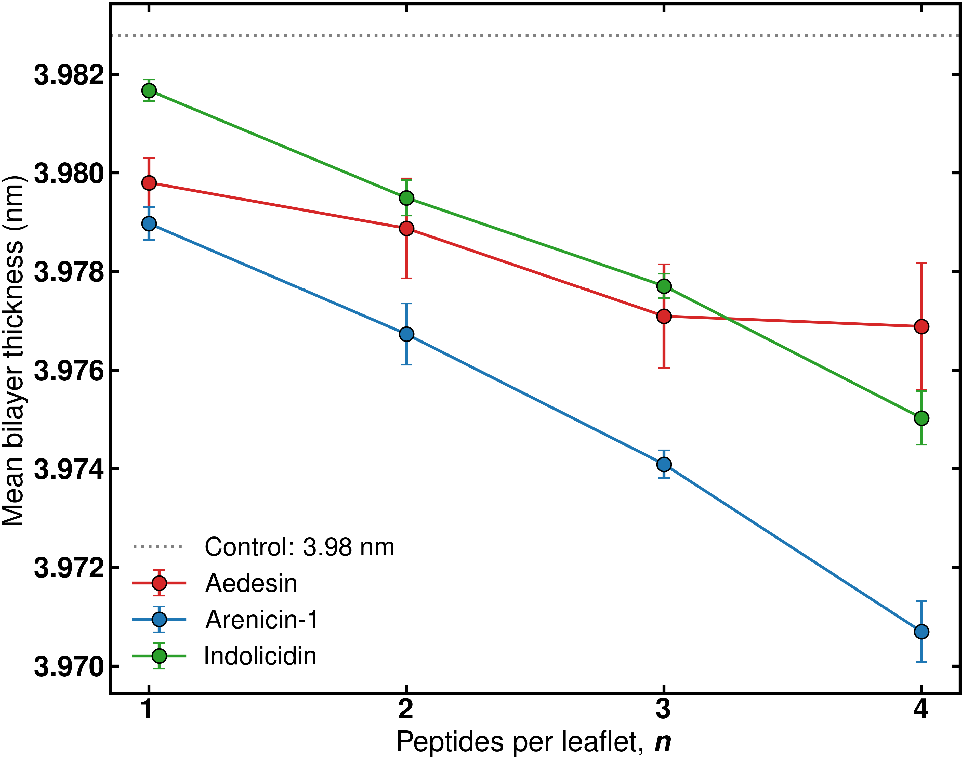
Mean bilayer thickness as a function of peptide number for the three AMP systems. Error bars are the standard error of the mean across replicas. The dashed gray line and shaded band show the lipid-only control. The ordering of bulk thinning (arenicin-1 > indolicidin > aedesin at n = 4) is the inverse of the K_*c*_ and K_*A*_ softening order, reflecting that the bulk geometric thickness probes direct PO_4_ displacement by peptides occupying the headgroup region rather than the wedge mechanical perturbation that dominates the elastic moduli.

**FIG. S6.**
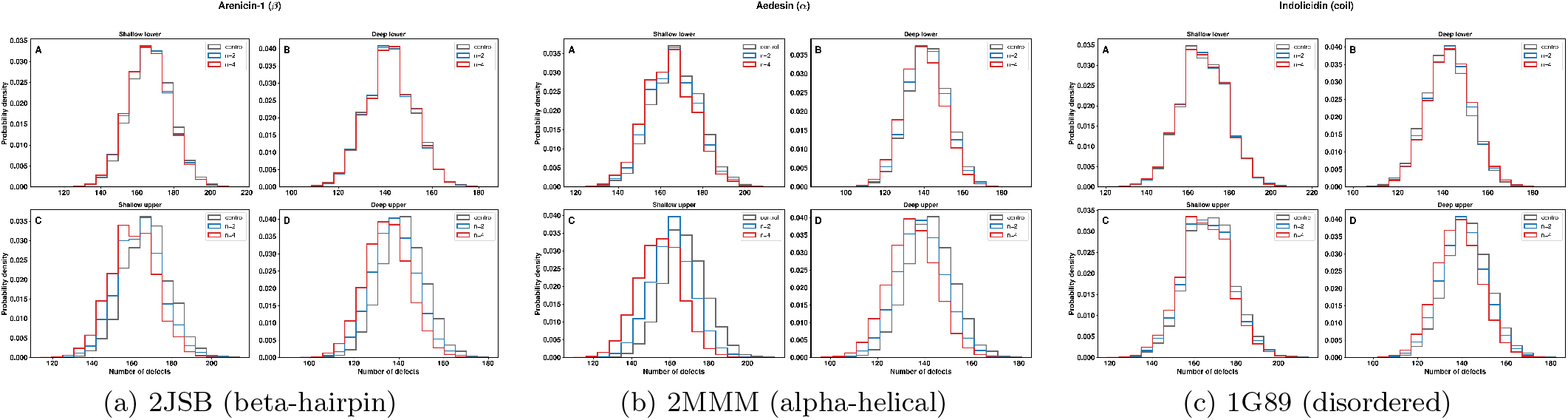
Per-frame distributions of the number of lipid packing defects for the three AMP systems at the highlighted concentrations (control vs n = 2 vs n = 4). Each sub-figure contains four panels: (A) shallow defects on the lower leaflet, (B) deep defects on the lower leaflet, (C) shallow defects on the upper (peptide facing) leaflet, and (D) deep defects on the upper leaflet. Histograms are density-normalized to enable direct comparison across conditions with different sample sizes. The upper leaflet deep defect panel (D) shows a systematic leftward shift of the defect-count distribution with increasing peptide loading for all three peptides. The upper leaflet shallow defect panel (C) shows a similar leftward shift for the structured peptides arenicin1 and aedesin but only a marginal shift for the disordered indolicidin. The lower leaflet panels (A, B) remain essentially superimposed across conditions for all three peptides.

**FIG. S7.**
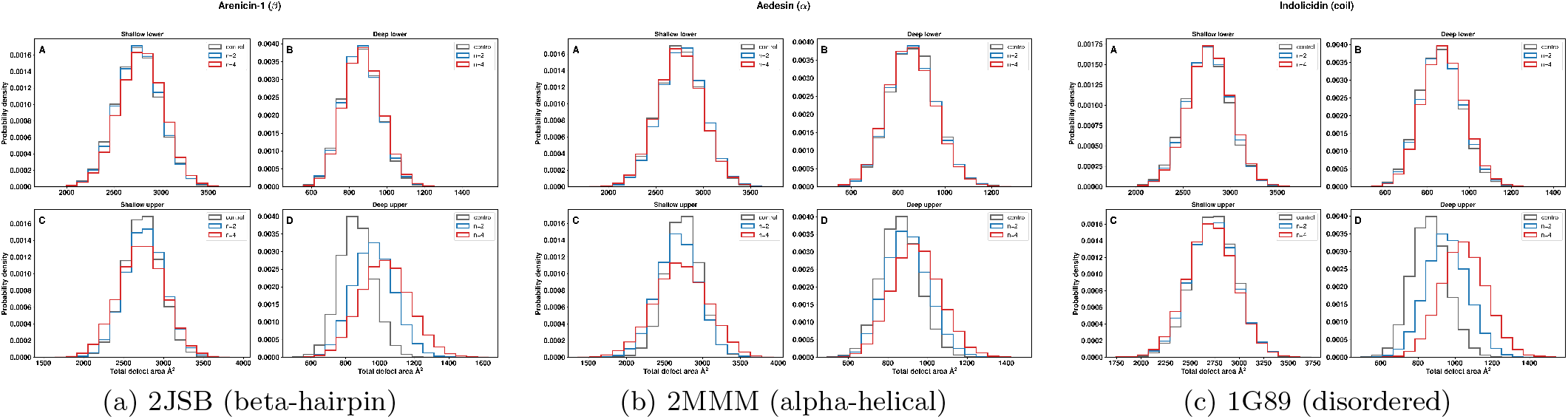
Per-frame distributions of the total lipid packing defect area for the three AMP systems at the highlighted concentrations (control vs n = 2 vs n = 4). Panel layout follows Figure S6: (A) shallow lower, (B) deep lower, (C) shallow upper, (D) deep upper. The deep-upper panels (D) show a clear rightward shift of the total-area distribution with peptide loading for all three peptides. The shallow-upper, shallow-lower, and deep-lower distributions remain essentially unchanged across conditions, consistent with the consolidation signature in the shallow channel (decreasing count, increasing individual defect area) being mass-conserving in integrated area.

