## Supplementary Information for "Dose-Dependent Softening of Bacterial Model Membranes by Structurally Distinct Antimicrobial Peptides: A Coarse-Grained Molecular Dynamics Study"

Roni Saiba, Krishnakanth Baratam, Debayan Chakraborty, and Satyavani Vemparala
*The Institute of Mathematical Sciences, C.I.T. Campus, Taramani, Chennai 600113, India and*
*Homi Bhabha National Institute, Training School Complex, Anushakti Nagar, Mumbai 400094, India*
(Dated: June 19, 2026)

The peptide-induced difference profiles $\Delta s(z) =$

$s_{\mathrm{peptide}}(z) - s_{\mathrm{control}}(z)$, shown in Figure S2, isolate the concentration dependent component of the stress redistribution and reveal a structural class dependent localization that matches the supraphosphate adsorption versus interfacial insertion framework developed in Section III C of the main text. For 1G89 (indolicidin, disordered), $\Delta s(z)$ develops a pronounced negative dip centered at $z \approx +1.5\,\mathrm{nm}$, slightly inboard of the $PO_4$ headgroup peak, with a magnitude that grows monotonically from $\sim -1000$ bar at n = 1 to $\sim -4000$ bar at n = 4. The inboard location of the dip is consistent with the per-bead z-distributions in Figure 3 of the main text, which place the tryptophan and arginine side chains of 1G89 substantially below the $PO_4$ plane. For 2JSB (arenicin-1, beta-hairpin), $\Delta s(z)$ shows a narrower and shallower dip localized at the $PO_4$ headgroup plane itself ($z \approx +2\,\mathrm{nm}$), reaching $\sim -1500$ bar at n = 4. The location and width of the 2JSB feature are consistent with the partial embedment of the compact beta-hairpin within the headgroup region (mean position 0.22 nm above $PO_4$). For 2MMM (aedesin, alpha-helical), the difference profile remains close to zero across all concentrations, with only small ($\sim \pm 100$ bar) fluctuations distributed across the upper leaflet. This is consistent with the 2MMM center of mass sitting 0.49 nm above the $PO_4$ plane, in the headgroup-water region, where the elastic network constrained alpha-helical rod displaces water rather than perturbing the chain packing stress directly.

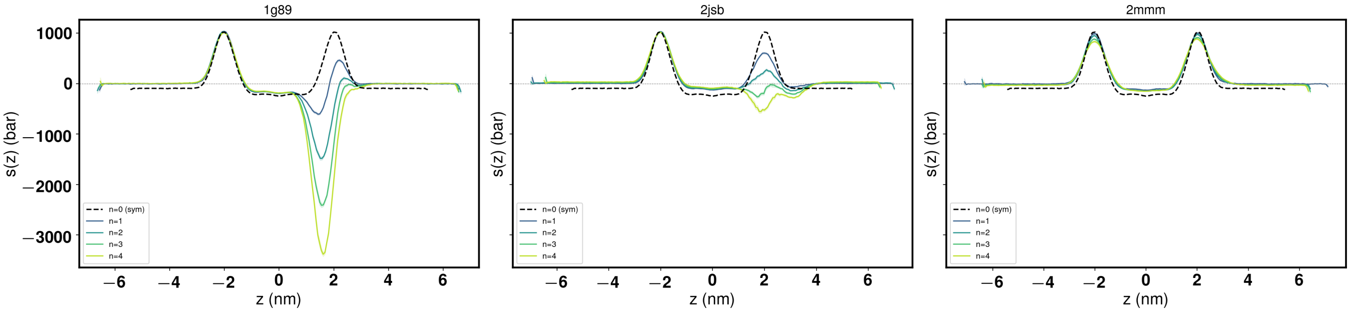


FIG. S1. Time-averaged lateral pressure profiles $s(z)$ for the three AMP systems at n = 1 to n = 4 peptides per leaflet. Solid colored lines are mean profiles across replicas (shaded bands: standard error of the mean); the dashed black line in each panel is the symmetrized lipid only control. The peptide binding leaflet is taken at $z > 0$ by convention; the distal leaflet at $z < 0$ remains essentially superposed on the control for all peptides and concentrations, demonstrating that the peptide induced stress perturbation is confined to the binding leaflet.

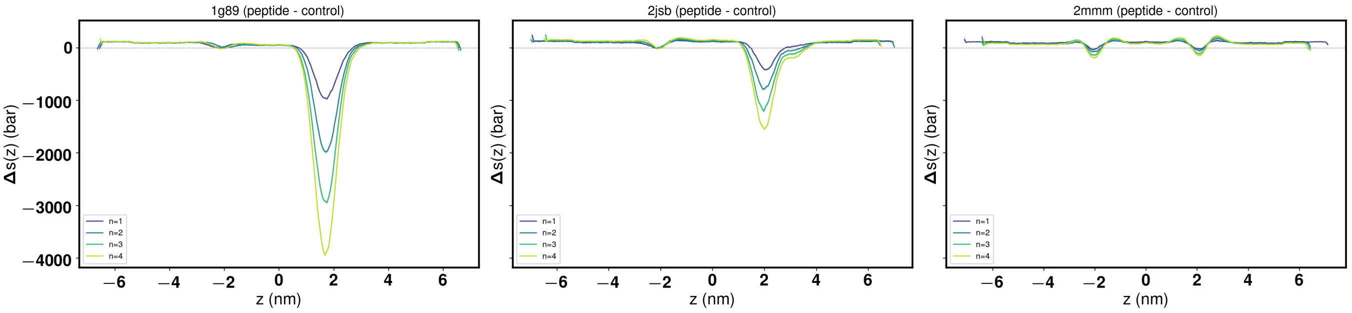


FIG. S2. Peptide-induced difference profiles $\Delta s(z) = s_{\text{peptide}}(z) - s_{\text{control}}(z)$ for the three AMP systems at n = 1 to n = 4. The structural class ordering recovers the supraphosphate adsorption versus interfacial insertion hierarchy of the main text: 1G89 (disordered) produces the deepest perturbation, localized inboard of the $PO_4$ plane consistent with side-chain insertion into the acyl chain region; 2JSB (beta-hairpin) produces an intermediate perturbation centered at the $PO_4$ plane consistent with partial headgroup embedment; 2MMM (alpha-helical) produces no resolvable perturbation within the nonbonded stress profile, consistent with its position 0.49 nm above the $PO_4$ plane in the headgroup-water region. The distal leaflet ($z < 0$) shows no peptide-induced perturbation for any system.

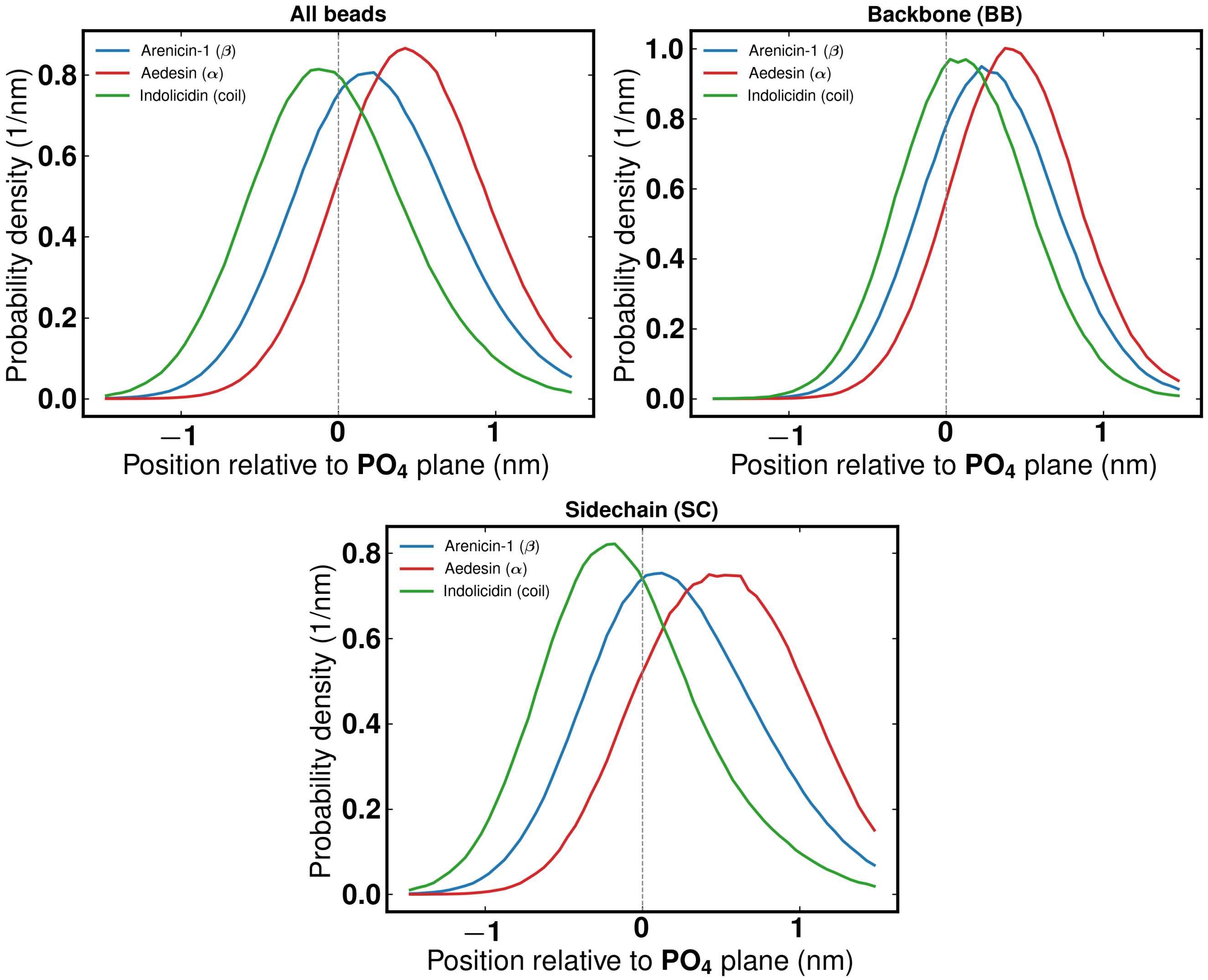


FIG. S3. Per-bead position distributions relative to the peptide facing $PO_4$ plane at n = 4 peptides per leaflet, for arenicin-1 (blue), aedesin (red), and idolicidin (green). The x-axis reports signed position in nm, with positive values indicating residence above $PO_4$ (headgroup-water region) and negative values indicating residence below $PO_4$ (toward the acyl chain region); the dashed vertical line marks the upper leaflet $PO_4$ reference plane. Left panel shows all coarse-grained beads pooled; center panel shows backbone beads (BB) only; right panel shows sidechain beads (SC) only. Distributions are normalized to unit area. Aedesin and arenicin-1 keep the large majority of their beads above the $PO_4$ plane (85% and 67% respectively), consistent with their supraphosphate adsorption mode. Indolicidin has 56% of its beads below $PO_4$, with the sidechain distribution (SC fraction below $PO_4$: 62%) extending substantially further into the membrane than the backbone (BB fraction: 39%), reflecting the interfacial insertion of its tryptophan and arginine residues while the backbone remains flush with the headgroup plane.

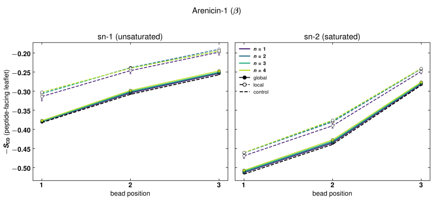


(a) Arenicin-1 (beta-hairpin)

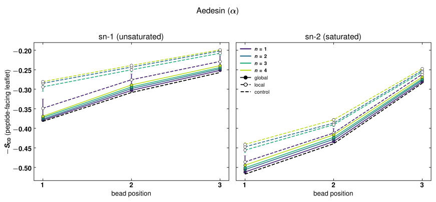


(b) Aedesin (alpha-helical)

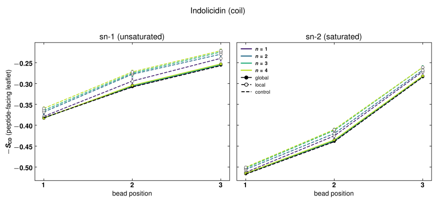


(c) Indolicidin (disordered)

FIG. S4. Per-chain SCD for each peptide class.

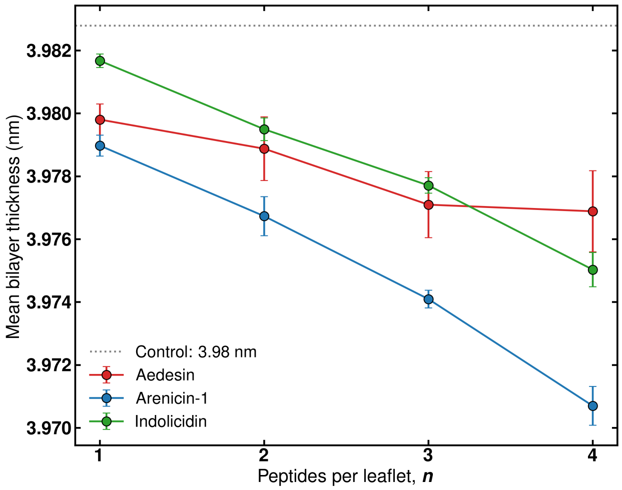


FIG. S5. Mean bilayer thickness as a function of peptide number for the three AMP systems. Error bars are the standard error of the mean across replicas. The dashed gray line and shaded band show the lipid-only control. The ordering of bulk thinning (arenicin-1 > indolicidin > aedesin at n = 4) is the inverse of the $K_c$ and $K_A$ softening order, reflecting that the bulk geometric thickness probes direct $PO_4$ displacement by peptides occupying the headgroup region rather than the wedge mechanical perturbation that dominates the elastic moduli.

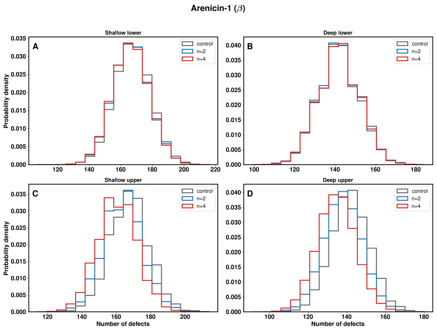


(a) 2JSB (beta-hairpin)

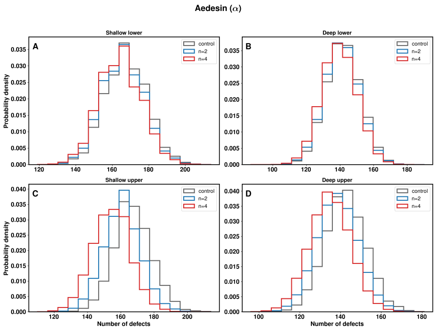


(b) 2MMM (alpha-helical)

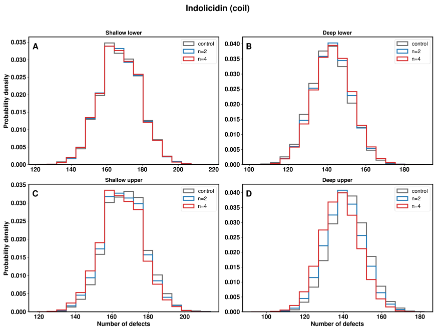


(c) 1G89 (disordered)

FIG. S6. Per-frame distributions of the number of lipid packing defects for the three AMP systems at the highlighted concentrations (control vs n = 2 vs n = 4). Each sub-figure contains four panels: (A) shallow defects on the lower leaflet, (B) deep defects on the lower leaflet, (C) shallow defects on the upper (peptide facing) leaflet, and (D) deep defects on the upper leaflet. Histograms are density-normalized to enable direct comparison across conditions with different sample sizes. The upper leaflet deep defect panel (D) shows a systematic leftward shift of the defect-count distribution with increasing peptide loading for all three peptides. The upper leaflet shallow defect panel (C) shows a similar leftward shift for the structured peptides arenicin1 and aedesin but only a marginal shift for the disordered indolicidin. The lower leaflet panels (A, B) remain essentially superimposed across conditions for all three peptides.

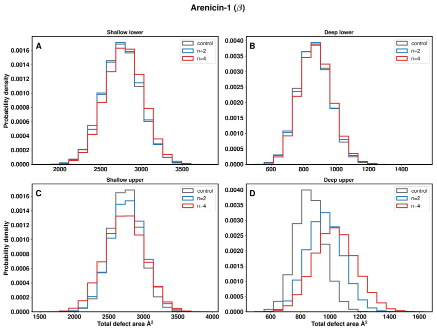


(a) 2JSB (beta-hairpin)

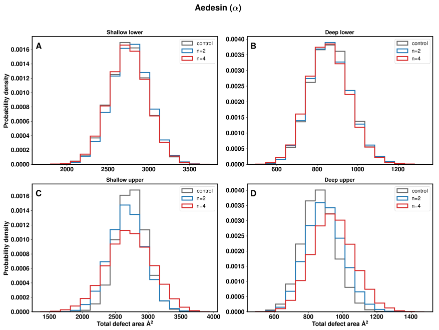


(b) 2MMM (alpha-helical)

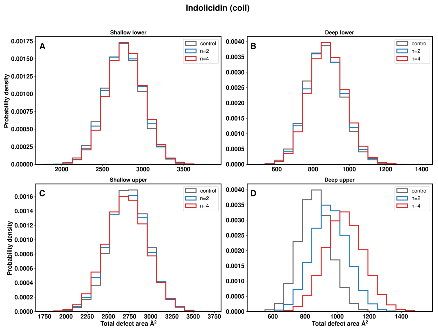


(c) 1G89 (disordered)

FIG. S7. Per-frame distributions of the total lipid packing defect area for the three AMP systems at the highlighted concentrations (control vs n = 2 vs n = 4). Panel layout follows Figure S6: (A) shallow lower, (B) deep lower, (C) shallow upper, (D) deep upper. The deep-upper panels (D) show a clear rightward shift of the total-area distribution with peptide loading for all three peptides. The shallow-upper, shallow-lower, and deep-lower distributions remain essentially unchanged across conditions, consistent with the consolidation signature in the shallow channel (decreasing count, increasing individual defect area) being mass-conserving in integrated area.
